# Formation of ultralong DH regions through genomic rearrangement

**DOI:** 10.1101/809764

**Authors:** Brevin A. Smider, Vaughn V. Smider

**Affiliations:** The Applied Biomedical Science Institute, San Diego, CA 92127; The Scripps Research Institute, La Jolla, CA 92037

## Abstract

Cow antibodies are very unusual in having exceptionally long CDR H3 regions. The genetic basis for this length largely derives from long heavy chain diversity (DH) regions, with a single “ultralong” DH, IGHD8-2, encoding over fifty amino acids. Most bovine IGHD regions are homologous but have several nucleotide repeating units that diversify their lengths. Genomically, most DH regions exist in three clusters that appear to have formed from DNA duplication events. The cluster containing IGHD8-2 underwent a rearrangement and deletion event in relation to the other clusters in the region corresponding to IGHD8-2, with possible fusion of two DH regions and expansion of short repeats to form the ultralong IGHD8-2 gene. Length heterogeneity within DH regions is a unique evolutionary genomic mechanism to create immune diversity, including formation of ultralong CDR H3 regions.

## Introduction

Adaptive immunity arose in vertebrates through the ability to somatically alter antigen receptor (antibody and T-cell receptor) genes to form diverse repertoires which are selected to bind and neutralize invading pathogens. A key component of this system is the ability to perform recombination of variable (V), diversity (D), and joining (J) gene segments through the process of V(D)J recombination[1–3]. A diversity of V, D, and J elements, along with imprecise joining at the V-D and D-J junctions enables different amino acids to be encoded in key paratopic regions which impact antigen binding.

The third complementary determining region of the heavy chain (CDR H3) is particularly important in antibody molecules as it contains the greatest diversity and also usually makes the most extensive contact with antigen. Long CDR H3 regions are often found in broadly neutralizing antibodies targeting human immunodeficiency (HIV), influenza, and polio viruses[4–8], and are also thought to be important in binding challenging antigens like G-protein coupled receptors and protease active sites[9, 10]. Thus, genetic mechanisms to form long CDR H3s may be very important in immune responses against key antigens.

In most organisms, the antibody CDR H3 forms a loop of 10-15 amino acids in length, and is encoded by the DH gene and associated recombinational junctions that form through VDJ recombination. Unusually long CDR H3s, such as those in broadly neutralizing anti-HIV antibodies, are often over 20 amino acids in length [4, 11–13]. The major determinants of CDR H3 length are the length of the germline encoded DH region, as well as somatic insertion of nucleotides (*e.g*. N- or P- nucleotides) at the V-D and D-J junctions. In humans, the longest DH region, IGHD3-16, encodes 12 amino acids.

Bovines are remarkable in having very long CDR H3 regions[14–24], with an average length of 26 amino acids [16] but with an exceptionally long subset of the repertoire (the “ultralong” CDR H3 antibodies) that can have CDR H3 lengths of up to seventy amino acids. These CDR H3 regions form their own independently folding mini domains comprised of a β-ribbon “stalk” that protrudes far from the typical paratope surface upon which sits a disulfide-bonded “knob”[21, 23, 25, 26]. Cows are the only species thus far investigated that can produce a broadly neutralizing antibody response against HIV, which is characterized by ultralong CDR H3 regions that penetrate the glycan shield of the spike protein to bind a conserved broadly neutralizing epitope in the CD4 binding region [27]. Cows are therefore unusual in producing long CDR H3s, and this unique repertoire has major functional relevance in neutralizing an antigen that is extremely challenging for repertoires of other mammalian species. Therefore, understanding the natural genetic and evolutionary mechanisms behind ultralong CDR H3 generation would be important in vaccine generation as well as therapeutic antibody discovery and development.

At least two evolutionary genetic events occurred which enabled formation of ultralong CDR H3 antibodies in cows. First, a unique VH region evolved as a result of an 8-basepair duplication at the 3’ end of IGHV1-7[16]. This particular variable region is the only VH region used in ultralong CDR H3 antibodies, and the short duplication directly encodes the ascending strand of the stalk region characteristic of these antibodies. Second, a very long DH region is found in cattle, IGHD8-2, which encodes 49 amino acids[23, 28–30]. Antibodies with ultralong CDR H3 regions invariably use IGHV1-7 and IGHD8-2[8, 16]. Here we examine the genetic features at work in the evolution of this unusually long DH region of cattle.

## Results

### DH cluster 2 has a significant deletion

We analyzed the DH regions of the recent assembly of the *Bos taurus* immunoglobulin heavy chain locus [29]for features associated with the ultralong IGHD8-2 region. Of particular note, the DH regions at the heavy chain locus are divided into “clusters” that arose from duplication events through evolution. TheIMGT naming nomenclature for DH regions includes numerical designations for the family and cluster of each gene; for example, IGHD3-2 is in family 3 and located in cluster 2 [16, 31–33]. There are four clusters, with clusters 2-4 being highly homologous with nucleotide identities of 92% (cluster 2 vs cluster 3), 99.7% (cluster 3 vs cluster 4), and 92% (cluster 2 vs cluster 4). The sequences of the DH regions located within the clusters are also highly homologous, with DH regions occupying analogous locations being 96% to 100% identical at the nucleotide level (Supplemental Figure 1). A major discrepancy in the cluster sequences, however, is that cluster 2 (3480 nucleotides) is 358 and 364 nucleotides shorter than clusters 3 (3838 nt) and 4 (3844 nt), respectfully. Additionally, cluster 2 is comprised of only five DH regions, with one of them being the ultralong IGHD8-2, whereas clusters 3 and 4 are comprised of six DH regions (Figure 1). Thus, cluster 2 appears to have a significant genomic deletion in relation to the highly homologous clusters 3 and 4. We hypothesized that this deletion might be related to formation of the ultralong IGHD8-2 region located in cluster 2. In simplistic terms, one explanation for formation of an ultralong DH region would be by fusion of two DH regions through deletion of intragenic sequence, with the fusion maintaining recombination signal sequences of each DH at both the 5’ and 3’ ends.

**Figure 1.**
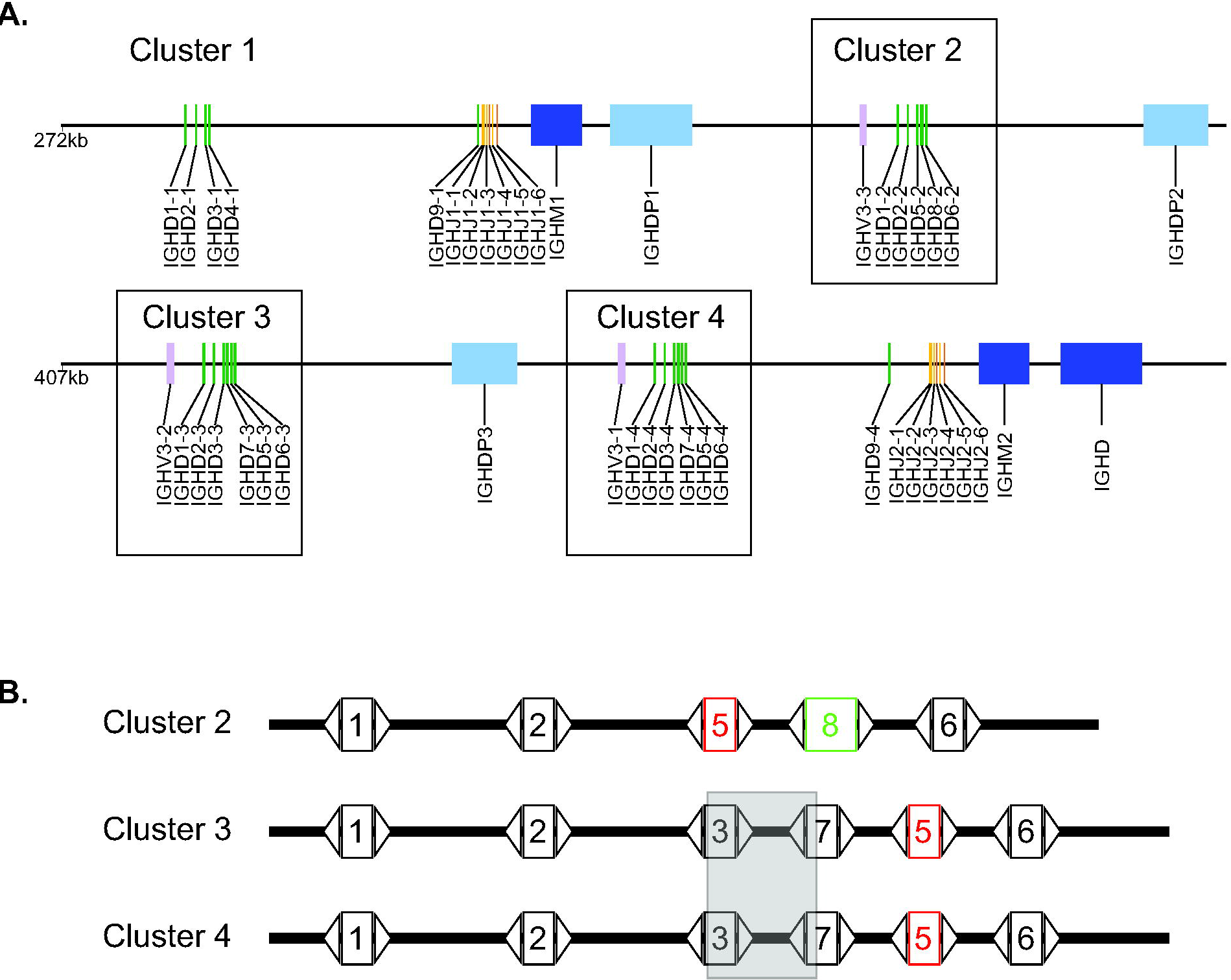
Schematic of D region clusters at the *Bos taurus* immunoglobulin heavy chain locus. (A) D-region cluster 2, comprising an ultralong IGHD, is shorter than highly homologous clusters. The DH regions are organized in four clustersat the immunoglobulin heavy chain locus on *Bos taurus* chromosome 21. Three clusters are highly homologous (clusters 2, 3 and 4 which are boxed). Green rectangles represent DH regions; orange, JH regions; blue, CH regions; light blue, pseudogene CH regions; and light pink, pseudogene VH regions. The entire locus is not shown; VH regions are upstream and remaining constant regions are downstream of the region shown. (B) Cluster 2 has a deletion and rearrangement in relation to clusters 3 and 4. Aligned schematic of the DH regions and their locations within the clusters. The numbers inside the boxes indicate the family members of each DH (*e.g*. on the first line, “1” represents IGHD1-2, and “1” on the second line represents IGHD1-3, etc.). IGHD5 is labeled in red to illustrate its unusual location in cluster 2 relative to clusters 3 and 4. The ultralong DH, IGHD8-2, is outlined in green. The transparent grey box encompassing IGHD3 and IGHD7 regions represents the approximate region of a large nucleotide deletion in cluster 2 relative to clusters 3 and 4. Triangles represent the recombination signal sequences (RSS) containing heptamer, 12 basepair spacer, and nonamer regions.

### Cluster 2 has a short chromosomal rearrangement

To evaluate the location of the deletion in cluster 2 relative to clusters 3 and 4, we performed a series of sequence alignments of the clusters, the DH regions, and the intergenic regions (between DH regions). Indeed, the deletion in cluster 2 in relation to clusters 3 and 4 occurred at IGHD8-2, however the deletion was also associated with a larger chromosomal rearrangement. In this regard, IGHD5-2 in cluster 2 appears to have replaced the paralog for IGHD3-3 (cluster 3) and IGHD3-4 (cluster 4)(Figure 1, Supplemental Figures 2-3). The IGHD5 homologs are immediately 5’ of the IGHD6 family members in clusters 3 and 4, however IGHD5-2 is situated immediately 3’ of IGHD2-2 and immediately 5’ of the ultralong IGHD8-2 region in cluster 2 (Figure 1). There is no IGHD3 family member in cluster 2 (Supplemental Figure 3), with the paralog of IGHD3-3 and IGHD3-4 either deleted or fused to the adjacent DH region, which would be a paralog of IGHD7-3 (cluster 3) or IGHD7-4 (cluster 4). Global alignments of the clusters show deleted nucleotides at IGHD8-2 as well as the position occupied by family 5 genes in clusters 3 and 4 (*e.g*. between IGHD7 and IGHD6). Alignments of the intergenic regions show that the intergenic region corresponding to the sequence between IGHD3-3 and IGHD7-3 in cluster 3 (or IGHD3-4 and IGHD7-4 in cluster 4) is deleted in cluster 2 (Supplemental Figure 4). While IGHD5-2 has been transposed to a location 3’ to IGHD8-2, the actual genetic material deleted clearly includes IGHD3 and its 3’ intergenic region. Thus, one possibility is that the ultralong IGHD8-2 region resulted from a deletion and associated fusion of the cluster 2 paralogs of IGHD3-3 and IGHD7-3. However, local sequence alignment reveals that the 5’ end of IGHD6-3 is 91.2% identical to IGHD8-2 (89.4% for IGHD6-2) over the first 85 nucleotides, whereas IGHD3-3 (and IGHD3-4) is only 80% over the first 62 residues (Supplemental Figure 7). Of note, IGHD6 family sequences share a cysteine in the same position as the conserved cysteine in IGHD8-2, which is highly conserved in deep sequenced ultralong CDR H3 antibodies, and forms a conserved disulfide bond at the base of the ultralong CDR H3 stalk [23, 26]. Thus, donation of an IGHD6 to the 5’ end of an IGHD7 through a recombinational or gene conversion process is a likely mechanism to produce IGHD8-2. Given the high homology of many of the DH regions and intergenic regions, we cannot definitively identify exact chromosomal breakpoints and cannot rule out that other events could have occurred in conjunction with the deletion event of the intragenic region between IGHD3 and IGHD7. For example, gene conversion could alternatively have occurred between IGHD6 and IGHD7 paralogs, or a deletion event followed by insertions of repeats into an IGHD7 paralog could have occurred. However, RSS analysis indicates that the 5’ RSS of IGHD8-2 shares identity with either IGHD3 or IGHD6 families (Table 1), thus a fusion between IGHD6 and IGHD7 or gene conversion of IGHD6 into IGHD3 followed by fusion to IGHD7 are likely mechanisms to produce the IGHD8-2 gene through a fusion event. The 3’ RSS of IGHD8-2 is identical to IGHD7 genes, and local alignments show homology between IGHD7 and IGHD8-2, suggesting that a primordial IGHD7 paralog from cluster 2 now forms the 3’ region of IGHD8-2.

**Table 1.**
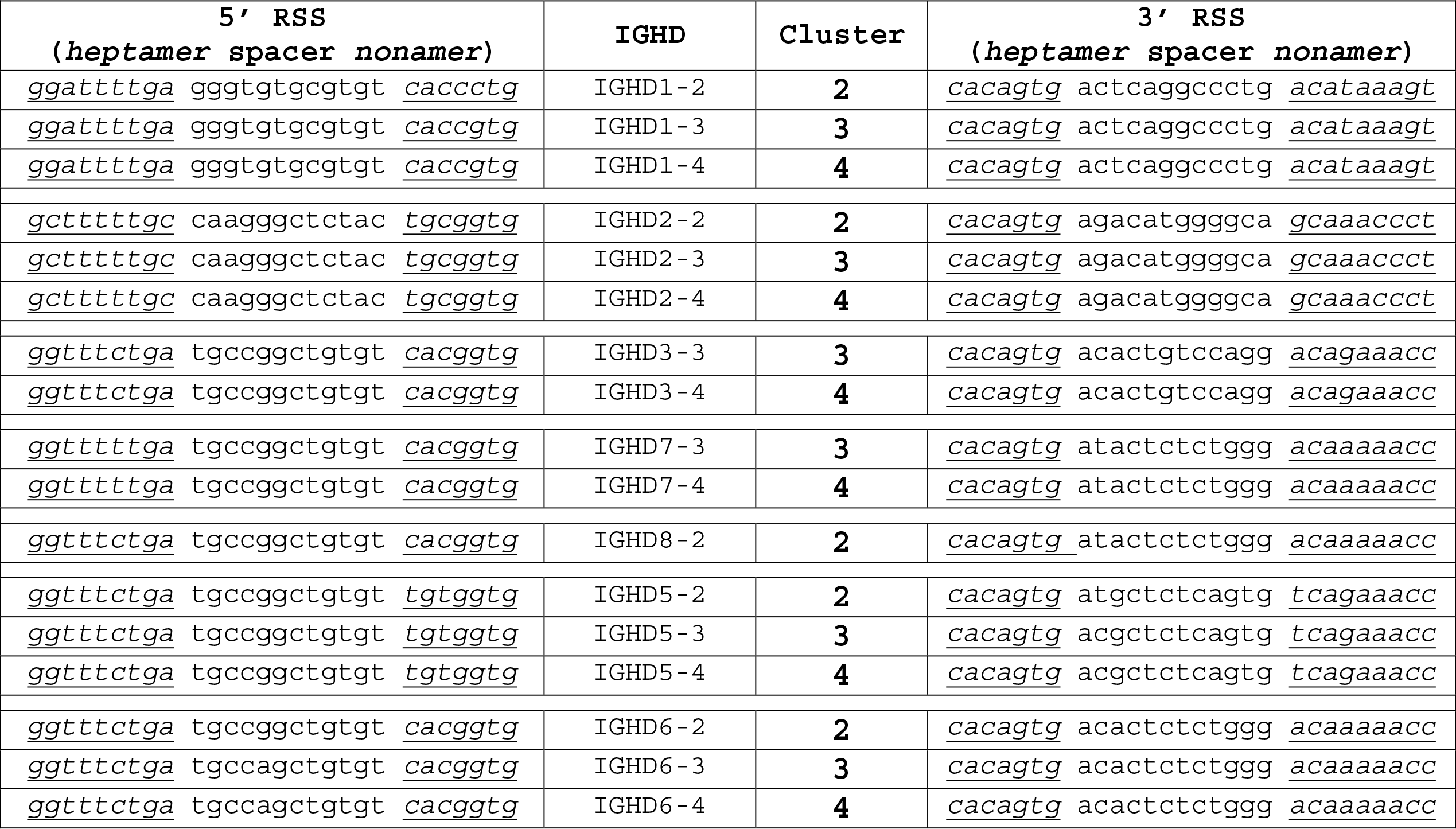
Recombination signal sequences of DH regions from clusters 2, 3 and 4.

### DH genes have expanded repeats

Bovine IGHD regions are comprised of multiple repeating short sequence motifs, with the major differences between several DH regions being length differences due to variable numbers of nucleotide repeats (Figure 2). IGHD7-4 is the second longest DH region, and only differs from IGHD7-3 (its paralog in cluster 3) by one repeat of TGGTTA, which results in a two amino acid change in length. IGHD7-3, IGHD7-4 and IGHD8-2 (the ultralong DH region) are very similar in having several repeating units, but with IGHD8-2 being dramatically longer. The 3’ ends of IGHD7-3 and IGHD7-4 are 85.6% and 77.4% identical to IGHD8-2 over the last 96 nucleotides, respectively (Supplementary Figure 5). The longer DH regions appear to be evolutionarily active in length evolution based on expanding or contracting repeats, as polymorphisms in IGHD8-2 *Bos taurus* differ in repeat lengths (Figure 2, Supplemental Figure 6). In this regard, two IGHD8-2 polymorphisms have been reported that differ in length and cysteine position, but share similar repeating nucleotide and amino acid sequences[29, 30, 34]. Related species like *Bos grunniens* (domestic Yak) and *Bison bison* (American buffalo) also have ultralong CDR H3 regions encoded by IGHD8-2 orthologs, but differ in their lengths due to apparent differences in hexanucleotide repeat expansion within the coding regions (Figure 2, Supplemental Figure 6). Thus, while two DH genes may have fused to form the long IGHD8-2 gene, nucleotide repeat expansion or contraction appears to also play a role in long DH region evolution in these species. To summarize, our analysis indicates that the most likely origin of IGHD8-2 is through a fusion event comprising the 5’ end of IGHD6 with the 3’ end of IGHD7 based on homology analysis, as well as the preservation of the codon encoding the nearly completely conserved first cysteine of the knob domain. This event, however, was associated with a larger chromosomal rearrangement that replaced an IGHD3 paralog and its 3’ intergenic region in cluster 2 with IGHD5-2 (Figure 3).

**Figure 2.**
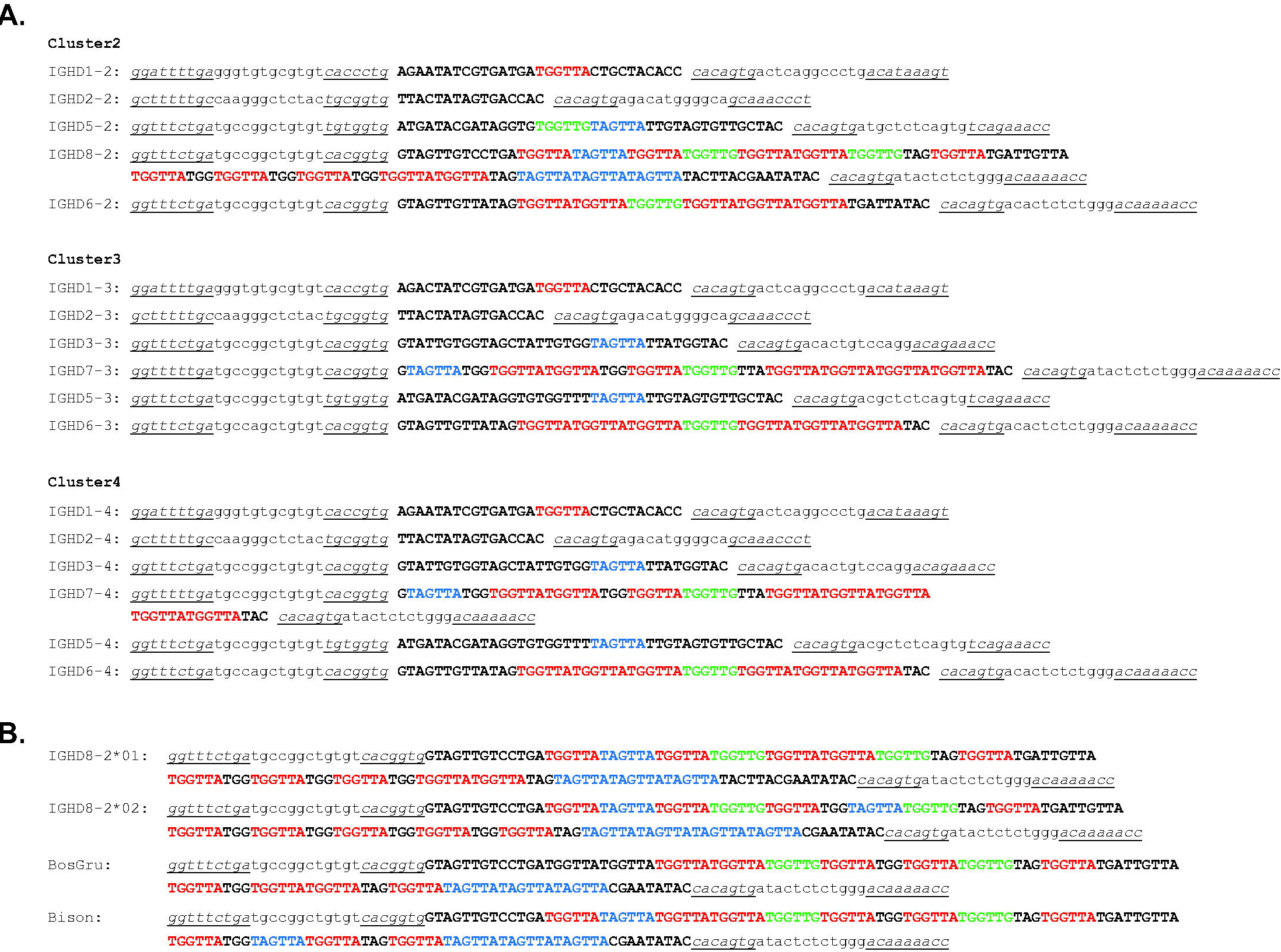
Bovine DH regions are characterized by repetitive sequences. (A) Nucleotide sequences of bovine DH regions in clusters 2-4. The RSS are in lowercase letters with the heptamer and nonamers in italics and underlined. The coding region of each DH is uppercase. Repeated sequences are colored red, blue and green. (B) Ungulate ultralong DH regions have different repeat lengths. The nucleotide sequences of polymorphic variants IGHD8-2*01 and IGHD8-2*02 for *Bos taurus* are compared to domestic yak (*Bos grunniens*; *BosGru*) and American bison (*Bison bison*; bison) orthologs of IGHD8-2.

**Figure 3.**
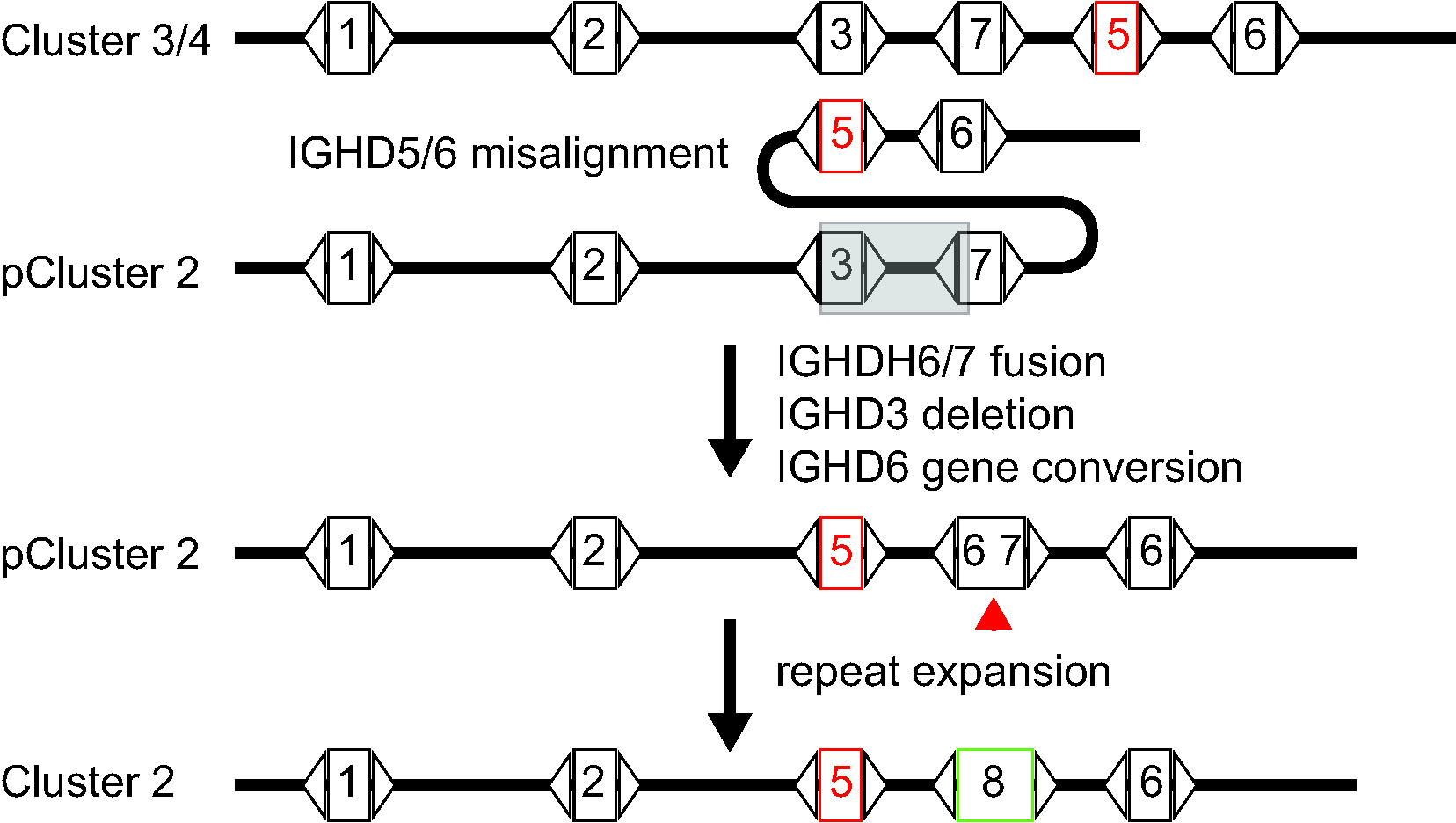
Model of deletion, fusion, and repeat expansion to form the ultralong DH region in cluster 2. Highly homologous sequences in clusters could misalign where IGHD5 and 6 pair with IGHD3 and 7 during replication processes. IGHD3 and its 3’ intergenic sequence is deleted (transparent grey rectangle). The short nucleotide repeats found in IGHD6 and 7 could cause mispairing and fusion, creating a fused DH-DH region of IGHD6 and IGHD7. A gene conversion or duplication event of IGHD6 would be required under this scenario. Continued repeat expansion produces IGHD8-2 homologs (red arrow). “pCluster” denotes a hypothetical precursor cluster in evolution.

### Conclusions and Discussion

The antibody repertoire is a defining evolutionary feature of vertebrates. V(D)J recombination and its associated junctional diversity account for vast potential diversity in antibody receptors on naïve B-cells, with an ability to bind with low affinity to most antigens. Most species utilize many V, D, and J gene segments which produce a great combinatorial potential at the heavy and light chain loci. Some species, however, have fewer functional V, D, and J segments and may use additional mechanisms to add diversity to their repertoires. Cows, in particular, have few VH and DH regions, but have very long CDR H3 regions. The homologous DH regions are cysteine rich, and diversity can be generated through both germline and somatically generated cysteines, which can form a diverse array of potential disulfide bonded loops[23, 35, 36]. In the knob region of ultralong CDR H3s, a diversity of disulfide bond patterns has been observed in several crystal structures [23, 25, 26], and mutations to and from cysteine have been confirmed through deep sequence analysis. Thus, novel cysteines encoded in the DH regions contributes to structural diversity in bovine antibodies.

The length of the DH regions in cows contributes to the overall increase in CDR H3 lengths in the antibody heavy chain repertoire. At the extreme, IGHD8-2 encodes 49 or 51 amino acids, depending on the polymorphic variant[29, 30, 34], and enables CDR H3 lengths of up to 70 amino acids in length. These CDR H3 regions form independently folding mini-domains comprised of a β-ribbon “stalk” and a disulfide-bonded “knob” that project far from the antigen surface. The sequence diversity of heavy chains with ultralong CDR H3 regions is enormous [16, 23], despite the fact that only one IGHD8-2 region (albeit with two polymorphic variants) is used in this entire class of antibodies. This vast diversity is explained by the fact that cattle utilize AID mediated somatic hypermutation in the pre-immune repertoire, as opposed to after antigen exposure as in other species, and this robust mutation induction substantially diversifies the repertoire through amino acid changes, cysteine mutations to alter disulfide loops, and a substantial proportion of deletional events which can also impact loop structures[16, 21].

All of these diversifying events use the germline IGHD regions as a template during repertoire formation. The IGHD templates are characterized by comprising multiple AID SH “hotspots” as well as nucleotide repeats that preferentially encode Ser, Gly, Tyr, and Cys, often in several repeating units like Gly-Tyr-Gly or Gly-Tyr-Ser. Here we show that nucleotide repeating units differ between IGHD paralogs derived from different clusters, and that the unusual ultralong IGHD8-2 region likely formed from a DH-DH fusion in cluster 2 of primordial IGHD6 (or IGHD3) to IGHD7 family members. Clearly a substantial deletion event occurred in the region now encoding IGHD8-2, which can be explained by deletion of the 3’ end of IGHD3, the intergenic region between IGHD3 and IGHD7, and the 5’ end of IGHD7. However, this event was also associated with a more substantial rearrangement that additionally replaced the IGHD3 paralog with IGHD5. Given that homologous variants of IGHD8-2 within *Bos taurus*, as well as in *Bos grunniens* and *Bison bison* differ in the length of IGHD8-2 through differences in the number of short repeats, it is likely that repeat expansion played a role in IGHD8-2 evolution either with a genetic fusion, or with massive expansion in the absence of the fusion of two DH regions. The ability of the genome to diversify IGHD region lengths through genomic rearrangement and repeat expansion provides a novel genetic mechanism for Darwinian diversification of the vertebrate immune system.

## Materials and Methods

The DNA sequence encoding the bovine antibody heavy chain locus (accession no. KT723008)[29] was downloaded from the IMGT server (http://www.imgt.org/). Clusters 2, 3 and 4 were defined by the beginning of the Family 1 gene RSS to the end of the Family 6 gene RSS. Sequences of IGHD regions, intergenic regions, and clusters were derived using the with Bioconductor program using the R statistical program language [37]. Multiple sequence alignments were done using Clustal Omega, WebPrank, or Muscle (https://www.ebi.ac.uk/services). Local sequence alignments were done using Matcher (https://www.ebi.ac.uk/Tools/psa/). The *Bison bison* and *Bos grunniens* IGHD8-2 sequences were identified by BLAST search at the ensembl genome server (www.ensembl.org) using the *Bos taurus* IGHD8-2 gene as query. *Bison bison* and *Bos grunniens* IGHD8-2 genes were found within the genomic sequences with accession numbers XM_010833706.1 (Bison) and CM016710.1 (Yak).

## Acknowledgements

This work was funded by NIH grants R01 GM105826 and R01 HD088400 to V.V.S. We are grateful for helpful conversations regarding this work with Ali Torkamani and Michael Criscitiello.

## Author Contributions

Both authors performed bioinformatic analysis and made the figures. V.V.S. wrote the manuscript with input from B.A.S.

---

**Supplemental Figure 1.** Alignment of *Bos taurus* DH regions.

**Supplemental Figure 2.** Alignment of DH clusters 2, 3, and 4.

**Supplemental Figure 3.** Phylogenetic of DH paralogs from clusters 2 and 3.

**Supplemental Figure 4.** Phylogenetic analysis of intergenic regions from clusters 2 and 3.

**Supplemental Figure 5.** Alignment of IGHD7 family members with IGHD8-2.

**Supplemental Figure 6.** Identification ultralong IGHD region homologs in *Bos taurus*, *Bos grunniens* and *Bison bison*.

**Supplemental Figure 7.** Alignment of IGHD3 and IGHD6 family members with IGHD8-2.

**Supplemental Figure 8.** Fusion analysis of IGHD members to form IGHD8-2 homologs.

