## Supplemental Fig. 1 for "Formation of ultralong DH regions through genomic rearrangement"

```
Supplemental Figure 1.  Alignment of all bovine DH regions in clusters 2-4 was performed using Clustal Omega.  (A) Percent identity matrix. (B) Sequence alignment.  (C) Phylogenetic tree.


A.
#
#
#  Percent Identity  Matrix - created by Clustal2.1
#
#
	          2-2    2-3      2-4     1-3     1-2      1-4    5-2     5-3     5-4     3-3     3-4     7-3     7-4     8-2     6-2     6-3     6-4
     1: IGHD2_2     100.00  100.00  100.00   36.11   37.04   40.74   52.78   52.78   52.78   48.61   48.61   50.70   50.70   52.78   50.70   49.30   49.30
     2: IGHD2_3     100.00  100.00  100.00   36.11   37.04   40.74   52.78   52.78   52.78   48.61   48.61   50.70   50.70   52.78   50.70   49.30   49.30
     3: IGHD2_4     100.00  100.00  100.00   36.11   37.04   40.74   52.78   52.78   52.78   48.61   48.61   50.70   50.70   52.78   50.70   49.30   49.30
     4: IGHD1_3      36.11   36.11   36.11  100.00   97.70   98.85   39.80   40.82   40.82   43.48   43.48   37.07   36.07   41.62   47.87   46.81   46.81
     5: IGHD1_2      37.04   37.04   37.04   97.70  100.00   98.85   55.17   55.17   55.17   60.00   60.00   55.88   55.88   49.41   60.00   58.00   58.00
     6: IGHD1_4      40.74   40.74   40.74   98.85   98.85  100.00   58.62   58.62   58.62   63.33   63.33   58.82   58.82   50.59   62.00   60.00   60.00
     7: IGHD5_2      52.78   52.78   52.78   39.80   55.17   58.62  100.00   97.96   97.96   71.43   71.43   74.16   73.68   72.45   78.08   76.71   76.71
     8: IGHD5_3      52.78   52.78   52.78   40.82   55.17   58.62   97.96  100.00  100.00   72.53   72.53   73.03   72.63   71.43   79.45   78.08   78.08
     9: IGHD5_4      52.78   52.78   52.78   40.82   55.17   58.62   97.96  100.00  100.00   72.53   72.53   73.03   72.63   71.43   79.45   78.08   78.08
    10: IGHD3_3      48.61   48.61   48.61   43.48   60.00   63.33   71.43   72.53   72.53  100.00  100.00   78.05   78.41   79.35   87.84   85.14   85.14
    11: IGHD3_4      48.61   48.61   48.61   43.48   60.00   63.33   71.43   72.53   72.53  100.00  100.00   78.05   78.41   79.35   87.84   85.14   85.14
    12: IGHD7_3      50.70   50.70   50.70   37.07   55.88   58.82   74.16   73.03   73.03   78.05   78.05  100.00  100.00   84.55   95.71   95.71   95.71
    13: IGHD7_4      50.70   50.70   50.70   36.07   55.88   58.82   73.68   72.63   72.63   78.41   78.41  100.00  100.00   84.50   95.71   95.71   95.71
    14: IGHD8_2      52.78   52.78   52.78   41.62   49.41   50.59   72.45   71.43   71.43   79.35   79.35   84.55   84.50  100.00   89.47   89.47   89.47
    15: IGHD6_2      50.70   50.70   50.70   47.87   60.00   62.00   78.08   79.45   79.45   87.84   87.84   95.71   95.71   89.47  100.00   96.49   96.49
    16: IGHD6_3      49.30   49.30   49.30   46.81   58.00   60.00   76.71   78.08   78.08   85.14   85.14   95.71   95.71   89.47   96.49  100.00  100.00
    17: IGHD6_4      49.30   49.30   49.30   46.81   58.00   60.00   76.71   78.08   78.08   85.14   85.14   95.71   95.71   89.47   96.49  100.00  100.00


B.

IGHD2_2      GCTTTTTGCCAAGGGCTCTACTGCGGT---------------------------------	27
IGHD2_3      GCTTTTTGCCAAGGGCTCTACTGCGGT---------------------------------	27
IGHD2_4      GCTTTTTGCCAAGGGCTCTACTGCGGT---------------------------------	27
IGHD1_3      GGATTTTGAGGGTGTGCGTGTCACCGTGAGACTATCGTGATGATGGTTACTG--------	52
IGHD1_2      GGATTTTGAGGGTGTGCGTGTCACCCTGAGAATATCGTGATGATGGTTACTG--------	52
IGHD1_4      GGATTTTGAGGGTGTGCGTGTCACCGTGAGAATATCGTGATGATGGTTACTG--------	52
IGHD5_2      GGTTTCTGATGCCGGCTGTGTTGTGGTGA-------------------------------	29
IGHD5_3      GGTTTCTGATGCCGGCTGTGTTGTGGTGA-------------------------------	29
IGHD5_4      GGTTTCTGATGCCGGCTGTGTTGTGGTGA-------------------------------	29
IGHD3_3      GGTTTCTGATGCCGGCTGTGTCACGGTGGT------------------------------	30
IGHD3_4      GGTTTCTGATGCCGGCTGTGTCACGGTGGT------------------------------	30
IGHD7_3      GGTTTTTGATGCCGGCTGTGTCACGG----------------------------------	26
IGHD7_4      GGTTTTTGATGCCGGCTGTGTCACGG----------------------------------	26
IGHD8_2      GGTTTCTGATGCCGGCTGTGTCACGGTGGT--AGTTGTCCTGATGGTTATAGTTATGGTT	58
IGHD6_2      GGTTTCTGATGCCGGCTGTGTCACGGTGGT--AGTTGTTATAGTGGTTATGGTTATGGTT	58
IGHD6_3      GGTTTCTGATGCCAGCTGTGTCACGGTGGT--AGTTGTTATAGTGGTTATGGTTATGGTT	58
IGHD6_4      GGTTTCTGATGCCAGCTGTGTCACGGTGGT--AGTTGTTATAGTGGTTATGGTTATGGTT	58
             *  ** **          *

IGHD2_2      ------------------------------------------------------------	27
IGHD2_3      ------------------------------------------------------------	27
IGHD2_4      ------------------------------------------------------------	27
IGHD1_3      ----------------CTACACCCACAGTGACTCAGGCCCTGACATAAAGTCT------G	90
IGHD1_2      ----------------CTACACCCACAGTGACTCAGGCCCTGACATAAAGT---------	87
IGHD1_4      ----------------CTACACCCACAGTGACTCAGGCCCTGACATAAAGT---------	87
IGHD5_2      ------------------------------------------------------------	29
IGHD5_3      ------------------------------------------------------------	29
IGHD5_4      ------------------------------------------------------------	29
IGHD3_3      ------------------------------------------------------------	30
IGHD3_4      ------------------------------------------------------------	30
IGHD7_3      -------------------------------------------TGGTAGTTATGGTGGTT	43
IGHD7_4      -------------------------------------------TGGTAGTTATGGTGGTT	43
IGHD8_2      ATGGTTGTGGTTATGGTTATGGTTGTAGTGGTTATGATTGTTATGGTTATGGTGGTTATG	118
IGHD6_2      GTGGTTATGGTT------------------------------------------------	70
IGHD6_3      ATGGTTGTGGTT------------------------------------------------	70
IGHD6_4      ATGGTTGTGGTT------------------------------------------------	70


IGHD2_2      -----------------------------------------GTTACTATAGTGACCACCA	46
IGHD2_3      -----------------------------------------GTTACTATAGTGACCACCA	46
IGHD2_4      -----------------------------------------GTTACTATAGTGACCACCA	46
IGHD1_3      ACCCGCACAC-AGGTGTGGAGCTGGCCAATGCATCCCCAGGGGCACTGGGCTCCCAAGCA	149
IGHD1_2      ------------------------------------------------------------	87
IGHD1_4      ------------------------------------------------------------	87
IGHD5_2      -----------------TGATACGATAGGTGTGGTTGTAGTTATTGTAGTGTTGCTACCA	72
IGHD5_3      -----------------TGATACGATAGGTGTGGTTTTAGTTATTGTAGTGTTGCTACCA	72
IGHD5_4      -----------------TGATACGATAGGTGTGGTTTTAGTTATTGTAGTGTTGCTACCA	72
IGHD3_3      ------------------------ATTGTGGTAGCTATTGTGGTAGTTATTATGGTACCA	66
IGHD3_4      ------------------------ATTGTGGTAGCTATTGTGGTAGTTATTATGGTACCA	66
IGHD7_3      ATGGTTATGGTGGTTATGGTTGTTATGGTTATGGT------TATGGTTATGGTTATACCA	97
IGHD7_4      ATGGTTATGGTGGTTATGGTTGTTATGGTTATGGTTATGGTTATGGTTATGGTTATACCA	103
IGHD8_2      GTGGTTATGGTGGTTATGGTTATAGTAGTTATAGTTATAGTTATACTTACGAATATACCA	178
IGHD6_2      ------------------------------------------ATGGTTATGATTATACCA	88
IGHD6_3      ------------------------------------------ATGGTTATGGTTATACCA	88
IGHD6_4      ------------------------------------------ATGGTTATGGTTATACCA	88


IGHD2_2      CAGTGAGACATGGGGCAGCAAACCCT	72
IGHD2_3      CAGTGAGACATGGGGCAGCAAACCCT	72
IGHD2_4      CAGTGAGACATGGGGCAGCAAACCCT	72
IGHD1_3      AGGTGCCTATCCCCCCAACTCGGGAC	175
IGHD1_2      --------------------------	87
IGHD1_4      --------------------------	87
IGHD5_2      CAGTGATGCTCTCAGTGTCAGAAACC	98
IGHD5_3      CAGTGACGCTCTCAGTGTCAGAAACC	98
IGHD5_4      CAGTGACGCTCTCAGTGTCAGAAACC	98
IGHD3_3      CAGTGACACTGTCCAGGACAGAAACC	92
IGHD3_4      CAGTGACACTGTCCAGGACAGAAACC	92
IGHD7_3      CAGTGATACTCTCTGGGACAAAAACC	123
IGHD7_4      CAGTGATACTCTCTGGGACAAAAACC	129
IGHD8_2      CAGTGATACTCTCTGGGACAAAAACC	204
IGHD6_2      CAGTGACACTCTCTGGGACAAAAACC	114
IGHD6_3      CAGTGACACTCTCTGGGACAAAAACC	114
IGHD6_4      CAGTGACACTCTCTGGGACAAAAACC	114
```

**C.**

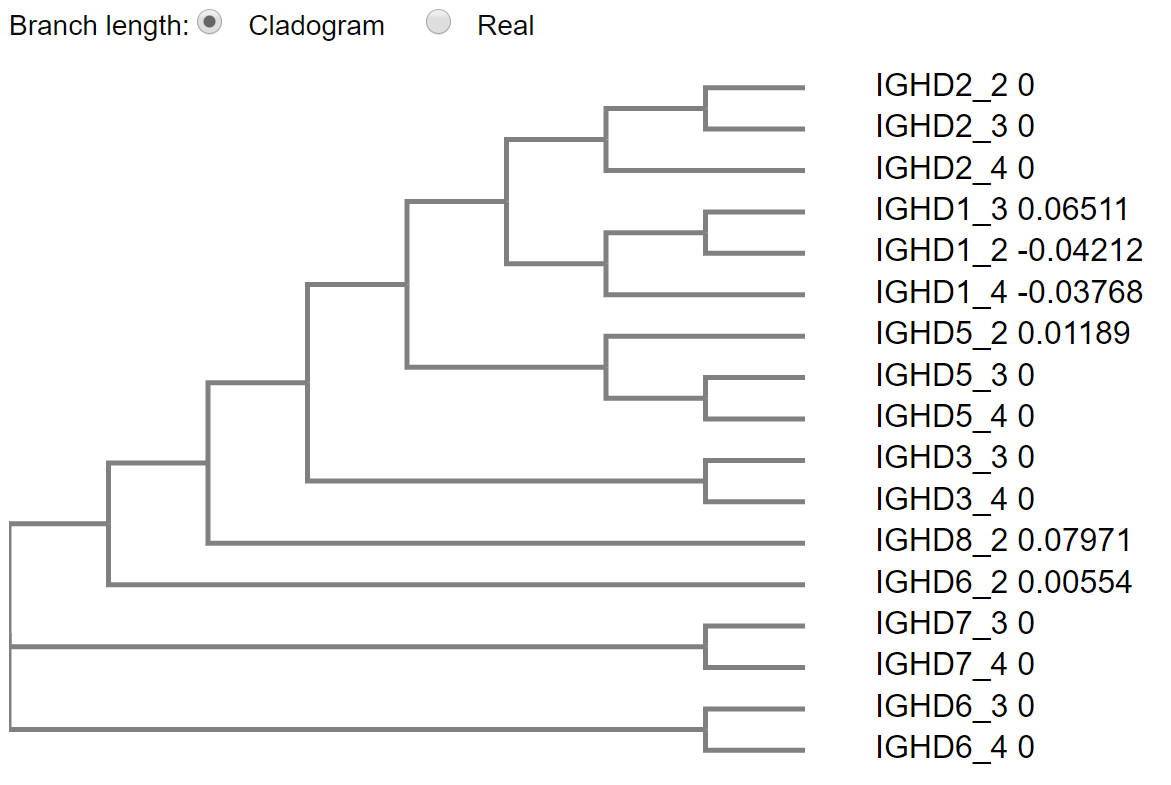
