## Supplemental Fig. 2 for "Formation of ultralong DH regions through genomic rearrangement"

**Supplemental Figure 2.** Global multiple sequence alignment of clusters 2-4 (above) using Webprank, or clusters 2 and 3 (below) using Stretcher. The DH regions are color coded as described in the Key. Note that IGHD5-2 is forced to align with IGHD3 in the global alignment. Supplemental Figures 3 and 4 show that the IGHD3 paralog does not exist (e.g. was deleted) in cluster 2.

```
Key
Green – Family 1
Light Green – Family 2
Red – Family 3
Purple – Family 7
Blue – IGHD8-2
Gold – Family 5
Orange – Family 6
Lowercase – RSS, with heptamer and nonamer in italics and underlined


#NEXUS
begin data;
dimensions ntax=3 nchar=3954;
format datatype=dna interleave=yes gap=-;
matrix

'Dcluster2'               ggattttgagggtgtgcgtgtcaccctgAGAATATCGTGATGATGGTTACTGCTACACCc
'Dcluster3'               ggattttgagggtgtgcgtgtcaccgtgAGACTATCGTGATGATGGTTACTGCTACACCc
'Dcluster4'               ggattttgagggtgtgcgtgtcaccgtgAGAATATCGTGATGATGGTTACTGCTACACCc

'Dcluster2'               acagtgactcaggccctgacataaagtCTGACCCGCACACAGGTGTGGAGCTTGCCAATG
'Dcluster3'               acagtgactcaggccctgacataaagtCTGACCCGCACACAGGTGTGGAGCTGGCCAATG
'Dcluster4'               acagtgactcaggccctgacataaagtCTGACCCGCACACAGGTGTGGAGCTGGCCAATG

'Dcluster2'               CATCCCCAGGGGCACTGGGCTCCCAAGCAAGGTGCCTATCCCCCCAACTCGGGACAGTGA
'Dcluster3'               CATCCCCAGGGGCACTGGGCTCCCAAGCAAGGTGCCTATCCCCCCAACTCGGGACAGTGA
'Dcluster4'               CATCCCCAGGGGCACTGGGCTCCCAAGCAAGGTGCCTATCCCCCCAACTCGGGACAGTGA

'Dcluster2'               GCCAGAAGGGATGTGTGGGCGTGGACCCTGGGCTCAGCCCACGGCCAGAGGACAGGGAAG
'Dcluster3'               GCCAGAAGGGATGTGTGGGCGTGGACCCTGGGCTCAGCCCACGGCCAGAGGACAGGGAAG
'Dcluster4'               GCCAGAAGGGATGTGTGGGCGTGGACCCTGGGCTCAGCCCACGGCCAGAGGACAGGGAAG

'Dcluster2'               GGCCCAGGGCTCCAGGCCACCTACCGAGAGCATTGAGCAAAGGTGGCTGAGGCCCCAATG
'Dcluster3'               GGCCCAGGGCTCCAGGCCACCTACCGAGAGCATTGAGCAAAGGTAGCTGAGGCCCCAATG
'Dcluster4'               GGCCCAGGGCTCCAGGCCACCTACCGAGAGCATTGAGCAAAGGTAGCTGAGGCCCCAATG

'Dcluster2'               AGGCAGCCATCCCAGGGAGCCCCTGACCCCTCCCCAGGACCAAGACTTGCCAGCAGGGCA
'Dcluster3'               AGGCAGCCATCCCAGGGAACCCCTGACCCCTCCCCAGGACCAAGACTTGCCAGCAGGGCA
'Dcluster4'               AGGCAGCCATCCCAGGGAACCCCTGACCCCTCCCCAGGACCAAGACTTGCCAGCAGGGCA

'Dcluster2'               GGGGTCCCGGCAAAGGGCCAGCCCTCCAGCCCTGGTGCACGCAC-CCCCTGCAGGCGAAG
'Dcluster3'               GGGGTCCCGGCAAAGGGCCAGCCCTCCAGCCCTGGTGCACACACCCCCCTGCAGGCGAAG
'Dcluster4'               GGGGTCCCGGCAAAGGGCCAGCCCTCCAGCCCTGGTGCACACACCCCCCTGCAGGCGAAG

'Dcluster2'               GGGGCACCTCTCGATTCCCCGGGGTGCTAGGGGGAGACAAGATCCAGGGGACTGTCCACA
'Dcluster3'               GGGGCACCTCTCGATTCCCCGGGGTGCTAGGGGGAGACAAGATCCAGGGGACTGTCCACA
'Dcluster4'               GGGGCACCTCTCGATTCCCCGGGGTGCTAGGGGGAGACAAGATCCAGGGGACTGTCCACA

'Dcluster2'               TTCCAGCAGGGACTCTGCTGGACAAGCCAGGGCTTCCAGGCCTGATCACAATCCATCACC
'Dcluster3'               TTCCAGCAGGGACTCTGCTGGACAAGCCAGGGCTTCCAGGCCTGATCACAACCCATCACC
'Dcluster4'               TTCCAGCAGGGACTCTGCTGGACAAGCCAGGGCTTCCAGGCCTGATCACAACCCATCACC

'Dcluster2'               CAAAATCAGCATGG-----GTGCCCAGGGGCCACTGAGGGCCTCAAGAAGGTGGGTCTCA
'Dcluster3'               CAAAATCAGCGTGGAGTGGGTGCCCAGGGGCCACTGAGGGCCTCAAGAAGGTGGGTCTCA
'Dcluster4'               CAAAATCAGCGTGGAGTGGGTGCCCAGGGGCCACTGAGGGCCTCAAGAAGGTGGGTCTCA

'Dcluster2'               ACCGTGCTCCATGGCCTGGAGCCGTGAGGGCACAGAGGGCATGAACCGACTGCAGAGGAG
'Dcluster3'               ACCGTGCTCCATGGCCTGGAGCCGTGAGGGCACAGAGGGCATGAACCGACTGCGGAGGAG
'Dcluster4'               ACCGTGCTCCATGGCCTGGAGCCGTGAGGGCACAGAGGGCATGAACCGACTGCGGAGGAG

'Dcluster2'               GGCCCTGCCCCTCAGCGAATGCCTCGTCCTGGGCCTCTGGGCCCCCGGGGTCAGGGCGGG
'Dcluster3'               GGCCCTGCCCCTCAGCGAATGCCTCGTCCTGGGCCTCTGGGCCCCCGGGGTCAGGGCGGG
'Dcluster4'               GGCCCTGCCCCTCAGCGAATGCCTCGTCCTGGGCCTCTGGGCCCCCGGGGTCAGGGCGGG

'Dcluster2'               GACGGGTGCAGGGTAGGGGGATGCAGCTCCCTGCACCTGCTCCCAGTCCCGCTGGGGCCA
'Dcluster3'               GACGGGTGCAGGGTAGGGGGATGCAGCTCCCTGCACCTGCTCCCAGTCCTGCTGGGGCCA
'Dcluster4'               GACGGGTGCAGGGTAGGGGGATGCAGCTCCCTGCACCTGCTCCCAGTCCTGCTGGGGCCA

'Dcluster2'               CGAGGATGAGCAGCTCAGCACTGCCCTCTGGGCTGGGCCAGCACAACTGGGCCCAGACTC
'Dcluster3'               CAAGGATGAGCAGCTCAGCACTGCCCTGTGGGCTGGGCCAGCACAACTGGGCCCAGACTC
'Dcluster4'               CAAGGATGAGCAGCTCAGCACTGCCCTGTGGGCTGGGCCAGCACAACTGGGCCCAGACTC

'Dcluster2'               CCAGGACGGGCAGCAGCAGGAATGACCCTGCTCTGCTCCTGGGTTTCCTGGCGGGAGGCG
'Dcluster3'               CCAGGACGGGCAGCAGCAGGAATGACCCTGCTCTGCTCCTGGGTTTCCTGGCGGGAGGCG
'Dcluster4'               CCAGGACGGGCAGCAGCAGGAATGACCCTGCTCTGCTCCTGGGTTTCCTGGCGGGAGGCG

'Dcluster2'               CCTACTGACCCCAGAGCACAACTGGCCTCAAGGTGACCGATACACAGTTTGTGGGCAATT
'Dcluster3'               CCTGCTGGCCCCAGAGCACAACCGGCCTCAAGGTGACCGATACACAGTTTGCAGGCAA-C
'Dcluster4'               CCTGCTGGCCCCAGAGCACAACCGGCCTCAAGGTGACCGATACACAGTTTGCAGGCAA-C

'Dcluster2'               TGGGTGCAATCTGTCAACCCAGGAGGCTGTCACTTGGCCAAGCCCAGCTTGGGGCCTCAG
'Dcluster3'               TGGGTAAAATCTGTCAACCCAGGAGGCTGTCACTTGGCTGAGCCCAGCTCGGGGCCTCAG
'Dcluster4'               TGGGTAAAATCTGTCAACCCAGGAGGCTGTCACTTGGCTGAGCCCAGCTCGGGGCCTCAG

'Dcluster2'               GAGAGCCAGCCTGGTCATCGTCCACCTGGCAGGAGACGCCTCCCACCGGCTCCAGCAGGT
'Dcluster3'               GAGAGCCAGCCTGGTCATCGTCCACCTGGCAGGAGATGCCTCCCAGGTGCTCCAGCAGGT
'Dcluster4'               GAGAGCCAGCCTGGTCATCGTCCACCTGGCAGGAGATGCCTCCCAGGTGCTCCAGCAGGT

'Dcluster2'               CGCCTGTGGCACCATGCCCTCCACAAGCCCTGAGCCCAGCACCAACCCACCCACAGCA--
'Dcluster3'               CTCCTGTGGCGCCATGCCCTCCACAAGCCCTGAGCCTGGCACCAACCCACCCACAGCCAG
'Dcluster4'               CTCCTGTGGCGCCATGCCCTCCACAAGCCCTGAGCCTGGCACCAACCCACCCACAGCCAG

'Dcluster2'               --CAGGCTGCCCAGCTCCCATGACTGTTCAGCCCTCTGAACGCCCCACATCCAGGGTCCC
'Dcluster3'               CCCAGGCCGCCCAGCCCCCATGACTGTTCAGCCCTCTGAACGCCCCACATCCAGGGTCCC
'Dcluster4'               CCCAGGCCGCCCAGCCCCCATGACTGTTCAGCCCTCTGAACGCCCCACATCCAGGGTCCC

'Dcluster2'               CACGCAGACGGCCTACAGCACGAACGGGgctttttgccaagggctctactgcggtgTTAC
'Dcluster3'               CACGCAGACGGCCTACAGCGCAAACGGGgctttttgccaagggctctactgcggtgTTAC
'Dcluster4'               CACGCAGACGGCCTACAGTGCAAACGGGgctttttgccaagggctctactgcggtgTTAC

'Dcluster2'               TATAGTGACCACcacagtgagacatggggcagcaaaccctGCCAACCGTATGGCCAACCA
'Dcluster3'               TATAGTGACCACcacagtgagacatggggcagcaaaccctGCCAACCGTATGGCCAACCA
'Dcluster4'               TATAGTGACCACcacagtgagacatggggcagcaaaccctGCCAACCGTATGGCCACCCA

'Dcluster2'               GCCCCCACGGCCGGGCCTCACGGCCTGAGCTTCAGAGACCAGACCTTCGGCCTTCTGGGC
'Dcluster3'               GCCCCCACGGCCGGGCCTCACGGCCTGAGCTTCAGAGACCAGACCTTCGGCCATCTGGGC
'Dcluster4'               GCCCCCACGGCCGGGCCTCACGGCCTGAGCTTCAGAGACCAGACCTTCGGCCATCTGGGC

'Dcluster2'               CACTCTGGGCGGATGTGGCCCGCGGGCCCGTATCCAGACTCTGAGATGACTGCACTGACC
'Dcluster3'               CACTCTGGGCGGATGCGGCCCACGGGCCCATATCCAGACTCTGAGATGACTGCACTGACC
'Dcluster4'               CACTCTGGGCGGATGCGGCCCATGGGCCCATATCCAGACTCTGAGATGACTGCACTGACC

'Dcluster2'               CTCCAGCCACACGTCCTAGCCCAGCAAGCTGGCCCTGACCACTAGCACTCAGTACACCCC
'Dcluster3'               CTCCAGCCACACATCCTAGCCCAGCAAGCTGGCCCTGACCACTAGCACTCAGTACACCCC
'Dcluster4'               CTCCAGCCACACATCCTAGCCCAGCAAGCTGGCCCTGACCACTAGCACTCAGTACACCCC

'Dcluster2'               CAAACAGGGTCAGGAAGGACAGAGCCCACGCTGGGCAAACCGTCAGGTCAAGTTTCCCAG
'Dcluster3'               CAAACAGGGTCAGGAAGGACAGAGCCCACGCTGGGCAAACCGTCAGGTCAAGTTTCCCAG
'Dcluster4'               CAAACAGGGTCAGGAAGGACAGAGCCCACGCTGGGCAAACCGTCAGGTCAAGTTTCCCAG

'Dcluster2'               GGGCAGGACCCGAGACACCTCCAACCCTGCTGGGAGCTCAGGGAAGCTCACAGCAGTGGA
'Dcluster3'               GGGCAGGACCTGAGACACCTCCAACCCTGCTGGGAGCTCAGGGAAGCTCACAGCAGTGGA
'Dcluster4'               GGGCAGGACCCGAGACACCTCCAACCCTGCTGGGAGCTCAGGGAAGCTCACAGCAGTGGA

'Dcluster2'               CGCTGGAGCTGGGCCTGTACTCAACTGCATAGGAGCTCACCAGGGAGCAGGCTGGGAGGG
'Dcluster3'               CGCTGGAGCTGGGCC-GTACTCAACTGCATAGGAGCTCGCCAGGGAGCAGGCTGGGAGGG
'Dcluster4'               TGCTGGAGCTGGGCCTGTACTCAACTGCATAGGAGCTCGCCAGGGAGCAGGCTGGGAGGG

'Dcluster2'               GTTCATGCAGAGGGAGCAGCTCCCAGCATGCTGCCAGACCCGAGGTAGGGGAGTCCGGAT
'Dcluster3'               GTTCATGCAGAGGCAGCAGCTCCCAGCATGCTGCCAGACCCGAGGTAGGGGAGTCCGGAT
'Dcluster4'               GTTCATGCAGAGGCAGCAGCTCCCAGCATGCTGCCAGACCCGAGGTAGGGGAGTCCGGAT

'Dcluster2'               CCAGGCCCGGCCTGAAGGGGCTCGGGGGGCCTCTCACATGTAGCAGGGGTGTGCAGGTCC
'Dcluster3'               CCAGGCCCGGCCTGAAGGGGCTCGGGGGGCCTCTCACATGTAGCAGGGGTGTGCAGGTCC
'Dcluster4'               CCAGGCCCGGCCTGAAGGGGCTCGAGGGGCCTCTCACATGTAGCAGGGGTGTGCAGGTCC

'Dcluster2'               CTTGTGCAGTGTCCAGAGATCATGGACAGGGCGGGCACCGCACCAGAAGCCAAGCCAGTG
'Dcluster3'               CTTGTGCAGTGTCCAGAGATCATGGACAGGGCGGGCACTGCACCAGAAGCCAAGCCAGTG
'Dcluster4'               CTTGTGCAGTGTCCAGAGATCATGGACAGGGCGGGCACTGCACCAGAAGCCAAGCCAGTG

'Dcluster2'               ACCAGCCTCTGCAGGAAGCACCCATGGGAGCCAACACCTAAGTCTCCCTCCCAGCCCTGC
'Dcluster3'               ACCAGCCTCTGCAGGAAGCGCCCGTGGGAGCCAACACCTAAGTCTCCCTCCCAGCCCTGC
'Dcluster4'               ACCAGCCTCTGCAGGAAGCGCCCGTGGGAGCCAACACCTAAGTCTCCCTCCCAGCCCTGC

'Dcluster2'               AGCGTGGCCACCCACCACCCCACAGGTCAGCCTGGGGAGCTCAAGACGGACTGGCGTCCC
'Dcluster3'               AGCGTGGCCACCCACCACCCCACAGGTCAGCCTGGGGAGCTCAAGACGGACTGGCGTCCC
'Dcluster4'               AGCGTGGCCACCCACCACCCCACAGGTCAGCCTGGGGAGCTCAAGACGGACTGGCGTCCC

'Dcluster2'               TACCTGCCAGGTTCCTGTGCAGGATGCTGTCCACGGCCGCAATGCTTTGGCCTCTGCTCA
'Dcluster3'               TACCTGCCAGGTTCCTGTGCAGGACGCTGTCCACAGGCGCAATGCTTGGGCCTCTGCTCA
'Dcluster4'               TACCTGCCAGGTTCCTGTGCAGGACGCTGTCCACAGGCGCAATGCTTGGGCCTCTGCTCA

'Dcluster2'               GCTTCTGAGGTTCTCCGGCCCACCACCACTGACCTGAATGGGCTTGGAGTGCAGGCCCAG
'Dcluster3'               GCTTCCGAGGTTCCCTGGCCCACCACCACCGACCTGAAGGGGCTCGGAGTGCAGGCCCAG
'Dcluster4'               GCTTCCGAGGTTCCCTGGCCCACCACCACCGACCTGAAGGGGCTCGGAGTGCAGGCCCAG

'Dcluster2'               CTGAACATCAGACCTGGGCTCAGAGAGCCACTCGGTCCAGGGTGCACTCAGCCGTGCACC
'Dcluster3'               CAGGACATAAGACCTGGGCTCAGATAGCCACTCGGTCCAGGGTGCACTCAGCCCTGCACC
'Dcluster4'               CAGGACATAAGACCTGGGCTCAGATAGCCACTCGGTCCAGGGTGCACTCAGCCCTGCACC

'Dcluster2'               TGGAGGAGGGTCCTCCATG-CCGCAGGGTTGGCTGGGCAAGGAGGCATCACAGACCACCC
'Dcluster3'               TGGAGGTGGGTCCTCCATGGCCGCAGCATTGGCTGGGCAAGGAGGCATCCCAGGCCACCC
'Dcluster4'               TGGAGGTGGGTCCTCCATGGCCGCAGCATTGGCTGGGCAAGGAGGCATCCCAGGCCACCC

'Dcluster2'               TGCTGTTGGGACGTCCAGCCCTGGACCAGGACTAGGTTCTCAGAATCCTCCCCACCACGG
'Dcluster3'               TGCCGTTGGGCCGTCCAGCCCTGGACCAGGACTAGGTTCTCAGAACCCTCCCCACCACGG
'Dcluster4'               TGCCGTTGGGCCGTCCAGCCCTGGACCAGGACTAGGTTCTCAGAACCCTCCCCACCACGG

'Dcluster2'               GACCTGTCCATACCTGGACCAGCTGACAGCTCCCAAGGGCTCCA-ACCCAGCAGACAAGA
'Dcluster3'               GACCCGTCCAGACCTGGGCCAGCTGACAGCTCCCAAGGGCTCCACACCCAGCAGACAAGA
'Dcluster4'               GACCCGTCCAGACCTGGGCCAGCTGACAGCTCCCAAGGGCTCCACACCCAGCAGACAAGA

'Dcluster2'               CCAA--GGGCGCTGAAGGCGAGCAGCTAGGgtttctgatgccggctgtgttgtggtgATG
'Dcluster3'               CCAAGGGGGCGCTGAAAACCAACAACTGGGgtttctgatgccggctgtgtcacggtgGTA
'Dcluster4'               CCAAGGGGGCGCTGAAAACCAACAACTGGGgtttctgatgccggctgtgtcacggtgGTA

'Dcluster2'               ATACGATAGGTGTGGTTGTAGTTATTGTAGTGTTGCTACcacagtgatgctctcagtgtc
'Dcluster3'               TTGTGGTAGCTATTGTGGTAGTTATTATGGT------ACcacagtgacactgtccaggac
'Dcluster4'               TTGTGGTAGCTATTGTGGTAGTTATTATGGT------ACcacagtgacactgtccaggac

'Dcluster2'               agaaaccCCTGCCCCGAGGTCCCAGACCAGGGAGCCTGTCCTCCAGGTACGGGGCACCGC
'Dcluster3'               agaaaccCCCGTCCCAAGGTCCCAGACCAGGGAGCCTGTCTTCCAAGACCGGGGTACCGC
'Dcluster4'               agaaaccCCCGTCCCAAGGTCCCAGACCAGGGAGCCTGTCTTCCAAGACCGGGGCACCGC

'Dcluster2'               CCTGCCCTCGTGACCATCAGACTGCACCCTGCACGCCCTCACCCTGCTGTCTCAGGGCAG
'Dcluster3'               CCTGCCCTCCTGACCATCAGACTGCACCCTGCACGCCCTCACCCCACTGGCTCAGGGAAG
'Dcluster4'               CCTGCCCTCCTGACCATCAGACTGCACCCTGCACGCCCTCACCCCACTGGCTCAGGGAAG

'Dcluster2'               CAACAGCCTCTGGAGGAGTCCTGGGCCAGGCACTCGGCTGTGGAGTCCCAGACAATGATA
'Dcluster3'               CAACAGCCTGAGGAAGGGTCCTGGGCCAGGCGCTGGGCTATGGGGACCCAGACAATGAGA
'Dcluster4'               CAACAGCCTGAGGAAGGGTCCTGGGCCAGGCGCTGGGCTATGGGGACCCAGACAATGAGA

'Dcluster2'               GCCTCAGCAGTTTCCAGCCACCTGACCCTGGAGGAGCTTGTGCAAGCTGGAGGCC-----
'Dcluster3'               CCCCCAGCAGGTTCCAGCCCCCTGACCCTGGAGGAACTT-------CCAGAGGCCTGTGC
'Dcluster4'               CCCCCAGCAGGTTCCAGCCCCCTGACCCTGGAGGAACTT-------CCAGAGGCCTGTGC

'Dcluster2'               --GGCTGGGGCCCGGAGCCTCAGAACCCTCCCTGCCAGGGGGCCCGTCCAGACCTGGACC
'Dcluster3'               CAGGCTGGGGACC-GACCCTAAGAACCCTCCCC-CGACAGGGCCCATCCAGA--------
'Dcluster4'               CAGGCTGGGGACC-GACCCTAAGAACCCTCCCC-CGACAGGGCCCATCCAGA--------

'Dcluster2'               AGCTGACAGCTCCCGAGGACTGCACGCCCACCAGACGAGACCAAGGGGGCGCTGAAGGCC
'Dcluster3'               ------CAGCTCTCGAGGTCTCCATGTCCACCAGACAAGACCAAGGGGGCGCTGAAGGCC
'Dcluster4'               ------CAGCTCTCGAGGTCTCCATGTCCACCAGACAAGACCAAGGGGGCGCTGAAGGCC

'Dcluster2'               AGCAGCTGGggtttctgatgccggctgtgtcacggtgGTAGTTGTCCTGATGGTTATAGT
'Dcluster3'               AGCAACTGGggtttttgatgccggctgtgtcacggtgGTAGTT------ATGGT------
'Dcluster4'               AGCAACTGGggtttttgatgccggctgtgtcacggtgGTAGTT------ATGGT------

'Dcluster2'               TATGGTTATGGTTGTGGTTATGGTTATGGTTGTAGTGGTTATGATTGTTATGGTTATGGT
'Dcluster3'               ---------------------GGTTATGGTTATGGTGGTTATGGTTGTT------ATGGT
'Dcluster4'               ---------------------GGTTATGGTTATGGTGGTTATGGTTGTTATGGTTATGGT

'Dcluster2'               GGTTATGGTGGTTATGGTGGTTATGGTTATAGTAGTTATAGTTATAGTTATACTTACGAA
'Dcluster3'               ---TATGGT------------TATGGTTATGGT---------------------------
'Dcluster4'               ---TATGGT------------TATGGTTATGGT---------------------------

'Dcluster2'               TATACcacagtgatactctctgggacaaaaaccCCTGCCCCTGAGGGTCCACGGCCAGGG
'Dcluster3'               TATACcacagtgatactctctgggacaaaaaccCCTGCCCCTGAGGGTCCACGACCAGGG
'Dcluster4'               TATACcacagtgatactctctgggacaaaaaccCCTGCCCCTGAGGGTCCACGACCAGGG

'Dcluster2'               ATCCTGGAGGCTGTGTTTGGGGCACCGCCCTGCCCTCCTGCCATTGGACTACAACCTGCA
'Dcluster3'               AATCTGTAAGCTGTGTTTGGGGCACCACCCTGCCCTCCTGCCATTGGACTGCACCCTGCA
'Dcluster4'               AATCTGTAAGCTGTGTTTGGGGCACCACCCTGCCCTCCTGCCATTGGACTGCACCCTGCA

'Dcluster2'               CGCCCTCACCCTGCTGACTCGGGCAGCAACAGCCTAAGATAAGAGTCCTGGGCCAGGCGC
'Dcluster3'               CGCCCTCACCCTGCTGACTCGGGCAGCAACAGCCTAAGATAAGGGTCCTGGGCCAGGAGC
'Dcluster4'               CGCCCTCACCCTGCTGACTCGGGCAGCAACAGCCTAAGATAAGGGTCCTGGGCCAGGAGC

'Dcluster2'               TGGGCTGTGGATACCCAGACAATGAGATCTCCAGCAGGTTCCCACCTCCTGACCCTGGAG
'Dcluster3'               TGGGCTGTGGATACCCAGACAATGAGATCTCCAGCAGGTTCCCGCCTCCTGATC-TGGAG
'Dcluster4'               TGGGCTGTGGATACCCAGACAATGAGATCTCCAGCAGGTTCCCGCCTCCTGATC-TGGAG

'Dcluster2'               GAACTTCCGCAGCCCTGTGGCAGGCTGGGGACCGACCCTAAGAACCCT-CCCCCCGC---
'Dcluster3'               GAACTGCCGCAGGCCTGTGCCAGGCTGGGGACCGACCCTAAGAACCCTCCCCCCCGCAGG
'Dcluster4'               GAACTGCCGCAGGCCTGTGCCAGGCTGGGAACCGACCCTAAGAACCCT-CCCCCCGCAGG

'Dcluster2'               ------------------------------------------------------------
'Dcluster3'               GCCATCCAGACCTGGGCCAGCTGACAGCTCCCGAGGACTCCACGTCCACCAGACGAGACC
'Dcluster4'               GCCATCCAGACCTGGGCCAGCTGACAGCTCCCGAGGACTCCACGTCCACCAGACGAGACC

'Dcluster2'               ------------------------------------------------------------
'Dcluster3'               AAGGGGACGCCGAAGGCCAGCAGCTGGggtttctgatgccggctgtgttgtggtgATGAT
'Dcluster4'               AAGGGGACGCCGAAGGCCAGCAGCTGGggtttctgatgccggctgtgttgtggtgATGAT

'Dcluster2'               ------------------------------------------------------------
'Dcluster3'               ACGATAGGTGTGGTTTTAGTTATTGTAGTGTTGCTACcacagtgacgctctcagtgtcag
'Dcluster4'               ACGATAGGTGTGGTTTTAGTTATTGTAGTGTTGCTACcacagtgacgctctcagtgtcag

'Dcluster2'               ------------------------------------------------------------
'Dcluster3'               aaaccCCTGCCCCGAGGTCCCAGACCAGGGAGCTTGTCCTCCAGGTCCGGGGTACGGCCC
'Dcluster4'               aaaccCCTGCCCCGAGGTCCCAGACCAGGGAGCTTGTCCTCCAGGTCCGGGGTATGGCCC

'Dcluster2'               ------------------------------------------------------------
'Dcluster3'               TGCCCTCCTGACCATCGGACTGCACCCTGCATGCCCTCACCCTGCTGACTCAGGGCAGCA
'Dcluster4'               TGCCCTCCTGACCATCGGACTGCACCCTGCATGCCCTCACCCTGCTGACTCAGGGCAGCA

'Dcluster2'               ------------------------------------------------------------
'Dcluster3'               ACAGAATGAGGAAGGGTCCTGGGCCAAGAGCTTGGCTGCTGGGACCCAGACAATGAGGCC
'Dcluster4'               ACAGAATGAGGAAGGGTCCTGGGCCAAGAGCTTGGCTGCTGGGACCCAGACAATGAGGCC

'Dcluster2'               ------------------------------------------------------------
'Dcluster3'               CCCAGCAAGTTCCAGCTCCCTGACCCTGGAGGAGGTTGTGCAGGTCAGAGGCCGGCTGGG
'Dcluster4'               CCCAGCAAGTTCCAGCCCCCTGACCCTGGAGGAGGTTGTGCAGGTCAGAGGCCGGCTGGG

'Dcluster2'               -----------------------------AGGGCCCGTCCAGACCTGAACCAGCTGACAG
'Dcluster3'               GACCGGATCCTCAGAACCCTCCCTGCCGCAGGGCCCGTCCAGA-CTGGACCAGCTGACAG
'Dcluster4'               GACCGGATCCTCAGAACCCTCCCTGCCGCAGGGCCCGTCCAGA-CTGGACCAGCTGACAG

'Dcluster2'               CTCCCGAGGGCTCCACGCCCCCCAGACGAGACCAAGGGGGCGCCGAAGGCCAGCAGCTGG
'Dcluster3'               CTCCCAAGGACTCCACGCCCACCAGACGAGACCAAGGGGGCGCCGAAGGCCAGCAGCCGG
'Dcluster4'               CTCCCAAGGACTCCACGCCCACCAGACGAGACCAAGGGGGCGCCGAAGGCCAGCAGCCGG

'Dcluster2'               ggtttctgatgccggctgtgtcacggtgGTAGTTGTTATAGTGGTTATGGTTATGGTTGT
'Dcluster3'               ggtttctgatgccagctgtgtcacggtgGTAGTTGTTATAGTGGTTATGGTTATGGTTAT
'Dcluster4'               ggtttctgatgccagctgtgtcacggtgGTAGTTGTTATAGTGGTTATGGTTATGGTTAT

'Dcluster2'               GGTTATGGTTATGGTTATGATTATACcacagtgacactctctgggacaaaaacc
'Dcluster3'               GGTTGTGGTTATGGTTATGGTTATACcacagtgacactctctgggacaaaaacc
'Dcluster4'               GGTTGTGGTTATGGTTATGGTTATACcacagtgacactctctgggacaaaaacc
;
end;

########################################
# Program: stretcher
# Rundate: Sat 17 Aug 2019 01:07:27
# Commandline: stretcher
#    -auto
#    -stdout
#    -asequence emboss_stretcher-I20190817-010723-0557-75507456-p2m.asequence
#    -bsequence emboss_stretcher-I20190817-010723-0557-75507456-p2m.bsequence
#    -datafile EDNAFULL
#    -gapopen 25
#    -gapextend 1
#    -aformat3 pair
#    -snucleotide1
#    -snucleotide2
# Align_format: pair
# Report_file: stdout
########################################

#=======================================
#
# Aligned_sequences: 2
# 1: Dcluster2
# 2: Dcluster3
# Matrix: EDNAFULL
# Gap_penalty: 25
# Extend_penalty: 1
#
# Length: 3947
# Identity:    3185/3947 (80.7%)
# Similarity:  3185/3947 (80.7%)
# Gaps:         574/3947 (14.5%)
# Score: 14191
#
#
#=======================================

Dcluster2          1 ggattttgagggtgtgcgtgtcaccctgAGAATATCGTGATGATGGTTAC     50
                     |||||||||||||||||||||||||.|||||.||||||||||||||||||
Dcluster3          1 ggattttgagggtgtgcgtgtcaccgtgAGACTATCGTGATGATGGTTAC     50

Dcluster2         51 TGCTACACCcacagtgactcaggccctgacataaagtCTGACCCGCACAC    100
                     ||||||||||||||||||||||||||||||||||||||||||||||||||
Dcluster3         51 TGCTACACCcacagtgactcaggccctgacataaagtCTGACCCGCACAC    100

Dcluster2        101 AGGTGTGGAGCTTGCCAATGCATCCCCAGGGGCACTGGGCTCCCAAGCAA    150
                     ||||||||||||.|||||||||||||||||||||||||||||||||||||
Dcluster3        101 AGGTGTGGAGCTGGCCAATGCATCCCCAGGGGCACTGGGCTCCCAAGCAA    150

Dcluster2        151 GGTGCCTATCCCCCCAACTCGGGACAGTGAGCCAGAAGGGATGTGTGGGC    200
                     ||||||||||||||||||||||||||||||||||||||||||||||||||
Dcluster3        151 GGTGCCTATCCCCCCAACTCGGGACAGTGAGCCAGAAGGGATGTGTGGGC    200

Dcluster2        201 GTGGACCCTGGGCTCAGCCCACGGCCAGAGGACAGGGAAGGGCCCAGGGC    250
                     ||||||||||||||||||||||||||||||||||||||||||||||||||
Dcluster3        201 GTGGACCCTGGGCTCAGCCCACGGCCAGAGGACAGGGAAGGGCCCAGGGC    250

Dcluster2        251 TCCAGGCCACCTACCGAGAGCATTGAGCAAAGGTGGCTGAGGCCCCAATG    300
                     ||||||||||||||||||||||||||||||||||.|||||||||||||||
Dcluster3        251 TCCAGGCCACCTACCGAGAGCATTGAGCAAAGGTAGCTGAGGCCCCAATG    300

Dcluster2        301 AGGCAGCCATCCCAGGGAGCCCCTGACCCCTCCCCAGGACCAAGACTTGC    350
                     ||||||||||||||||||.|||||||||||||||||||||||||||||||
Dcluster3        301 AGGCAGCCATCCCAGGGAACCCCTGACCCCTCCCCAGGACCAAGACTTGC    350

Dcluster2        351 CAGCAGGGCAGGGGTCCCGGCAAAGGGCCAGCCCTCCAGCCCTGGTGCAC    400
                     ||||||||||||||||||||||||||||||||||||||||||||||||||
Dcluster3        351 CAGCAGGGCAGGGGTCCCGGCAAAGGGCCAGCCCTCCAGCCCTGGTGCAC    400

Dcluster2        401 GCACCCCC-TGCAGGCGAAGGGGGCACCTCTCGATTCCCCGGGGTGCTAG    449
                     .||||||| |||||||||||||||||||||||||||||||||||||||||
Dcluster3        401 ACACCCCCCTGCAGGCGAAGGGGGCACCTCTCGATTCCCCGGGGTGCTAG    450

Dcluster2        450 GGGGAGACAAGATCCAGGGGACTGTCCACATTCCAGCAGGGACTCTGCTG    499
                     ||||||||||||||||||||||||||||||||||||||||||||||||||
Dcluster3        451 GGGGAGACAAGATCCAGGGGACTGTCCACATTCCAGCAGGGACTCTGCTG    500

Dcluster2        500 GACAAGCCAGGGCTTCCAGGCCTGATCACAATCCATCACCCAAAATCAGC    549
                     |||||||||||||||||||||||||||||||.||||||||||||||||||
Dcluster3        501 GACAAGCCAGGGCTTCCAGGCCTGATCACAACCCATCACCCAAAATCAGC    550

Dcluster2        550 ATGG-----GTGCCCAGGGGCCACTGAGGGCCTCAAGAAGGTGGGTCTCA    594
                     .|||     |||||||||||||||||||||||||||||||||||||||||
Dcluster3        551 GTGGAGTGGGTGCCCAGGGGCCACTGAGGGCCTCAAGAAGGTGGGTCTCA    600

Dcluster2        595 ACCGTGCTCCATGGCCTGGAGCCGTGAGGGCACAGAGGGCATGAACCGAC    644
                     ||||||||||||||||||||||||||||||||||||||||||||||||||
Dcluster3        601 ACCGTGCTCCATGGCCTGGAGCCGTGAGGGCACAGAGGGCATGAACCGAC    650

Dcluster2        645 TGCAGAGGAGGGCCCTGCCCCTCAGCGAATGCCTCGTCCTGGGCCTCTGG    694
                     |||.||||||||||||||||||||||||||||||||||||||||||||||
Dcluster3        651 TGCGGAGGAGGGCCCTGCCCCTCAGCGAATGCCTCGTCCTGGGCCTCTGG    700

Dcluster2        695 GCCCCCGGGGTCAGGGCGGGGACGGGTGCAGGGTAGGGGGATGCAGCTCC    744
                     ||||||||||||||||||||||||||||||||||||||||||||||||||
Dcluster3        701 GCCCCCGGGGTCAGGGCGGGGACGGGTGCAGGGTAGGGGGATGCAGCTCC    750

Dcluster2        745 CTGCACCTGCTCCCAGTCCCGCTGGGGCCACGAGGATGAGCAGCTCAGCA    794
                     |||||||||||||||||||.|||||||||||.||||||||||||||||||
Dcluster3        751 CTGCACCTGCTCCCAGTCCTGCTGGGGCCACAAGGATGAGCAGCTCAGCA    800

Dcluster2        795 CTGCCCTCTGGGCTGGGCCAGCACAACTGGGCCCAGACTCCCAGGACGGG    844
                     |||||||.||||||||||||||||||||||||||||||||||||||||||
Dcluster3        801 CTGCCCTGTGGGCTGGGCCAGCACAACTGGGCCCAGACTCCCAGGACGGG    850

Dcluster2        845 CAGCAGCAGGAATGACCCTGCTCTGCTCCTGGGTTTCCTGGCGGGAGGCG    894
                     ||||||||||||||||||||||||||||||||||||||||||||||||||
Dcluster3        851 CAGCAGCAGGAATGACCCTGCTCTGCTCCTGGGTTTCCTGGCGGGAGGCG    900

Dcluster2        895 CCTACTGACCCCAGAGCACAACTGGCCTCAAGGTGACCGATACACAGTTT    944
                     |||.|||.||||||||||||||.|||||||||||||||||||||||||||
Dcluster3        901 CCTGCTGGCCCCAGAGCACAACCGGCCTCAAGGTGACCGATACACAGTTT    950

Dcluster2        945 GTGGGCAATTTGGGTGCAATCTGTCAACCCAGGAGGCTGTCACTTGGCCA    994
                     |..|||||.| ||||..|||||||||||||||||||||||||||||||..
Dcluster3        951 GCAGGCAACT-GGGTAAAATCTGTCAACCCAGGAGGCTGTCACTTGGCTG    999

Dcluster2        995 AGCCCAGCTTGGGGCCTCAGGAGAGCCAGCCTGGTCATCGTCCACCTGGC   1044
                     |||||||||.||||||||||||||||||||||||||||||||||||||||
Dcluster3       1000 AGCCCAGCTCGGGGCCTCAGGAGAGCCAGCCTGGTCATCGTCCACCTGGC   1049

Dcluster2       1045 AGGAGACGCCTCCCACCGGCTCCAGCAGGTCGCCTGTGGCACCATGCCCT   1094
                     ||||||.||||||||...|||||||||||||.||||||||.|||||||||
Dcluster3       1050 AGGAGATGCCTCCCAGGTGCTCCAGCAGGTCTCCTGTGGCGCCATGCCCT   1099

Dcluster2       1095 CCACAAGCCCTGAGCCCAGCACCAACCCACCCACAGCA----CAGGCTGC   1140
                     ||||||||||||||||..|||||||||||||||||||.    |||||.||
Dcluster3       1100 CCACAAGCCCTGAGCCTGGCACCAACCCACCCACAGCCAGCCCAGGCCGC   1149

Dcluster2       1141 CCAGCTCCCATGACTGTTCAGCCCTCTGAACGCCCCACATCCAGGGTCCC   1190
                     |||||.||||||||||||||||||||||||||||||||||||||||||||
Dcluster3       1150 CCAGCCCCCATGACTGTTCAGCCCTCTGAACGCCCCACATCCAGGGTCCC   1199

Dcluster2       1191 CACGCAGACGGCCTACAGCACGAACGGGgctttttgccaagggctctact   1240
                     |||||||||||||||||||.|.||||||||||||||||||||||||||||
Dcluster3       1200 CACGCAGACGGCCTACAGCGCAAACGGGgctttttgccaagggctctact   1249

Dcluster2       1241 gcggtgTTACTATAGTGACCACcacagtgagacatggggcagcaaaccct   1290
                     ||||||||||||||||||||||||||||||||||||||||||||||||||
Dcluster3       1250 gcggtgTTACTATAGTGACCACcacagtgagacatggggcagcaaaccct   1299

Dcluster2       1291 GCCAACCGTATGGCCAACCAGCCCCCACGGCCGGGCCTCACGGCCTGAGC   1340
                     ||||||||||||||||||||||||||||||||||||||||||||||||||
Dcluster3       1300 GCCAACCGTATGGCCAACCAGCCCCCACGGCCGGGCCTCACGGCCTGAGC   1349

Dcluster2       1341 TTCAGAGACCAGACCTTCGGCCTTCTGGGCCACTCTGGGCGGATGTGGCC   1390
                     ||||||||||||||||||||||.||||||||||||||||||||||.||||
Dcluster3       1350 TTCAGAGACCAGACCTTCGGCCATCTGGGCCACTCTGGGCGGATGCGGCC   1399

Dcluster2       1391 CGCGGGCCCGTATCCAGACTCTGAGATGACTGCACTGACCCTCCAGCCAC   1440
                     |.|||||||.||||||||||||||||||||||||||||||||||||||||
Dcluster3       1400 CACGGGCCCATATCCAGACTCTGAGATGACTGCACTGACCCTCCAGCCAC   1449

Dcluster2       1441 ACGTCCTAGCCCAGCAAGCTGGCCCTGACCACTAGCACTCAGTACACCCC   1490
                     ||.|||||||||||||||||||||||||||||||||||||||||||||||
Dcluster3       1450 ACATCCTAGCCCAGCAAGCTGGCCCTGACCACTAGCACTCAGTACACCCC   1499

Dcluster2       1491 CAAACAGGGTCAGGAAGGACAGAGCCCACGCTGGGCAAACCGTCAGGTCA   1540
                     ||||||||||||||||||||||||||||||||||||||||||||||||||
Dcluster3       1500 CAAACAGGGTCAGGAAGGACAGAGCCCACGCTGGGCAAACCGTCAGGTCA   1549

Dcluster2       1541 AGTTTCCCAGGGGCAGGACCCGAGACACCTCCAACCCTGCTGGGAGCTCA   1590
                     ||||||||||||||||||||.|||||||||||||||||||||||||||||
Dcluster3       1550 AGTTTCCCAGGGGCAGGACCTGAGACACCTCCAACCCTGCTGGGAGCTCA   1599

Dcluster2       1591 GGGAAGCTCACAGCAGTGGACGCTGGAGCTGGGCCTGTACTCAACTGCAT   1640
                     ||||||||||||||||||||||||||||||||||| ||||||||||||||
Dcluster3       1600 GGGAAGCTCACAGCAGTGGACGCTGGAGCTGGGCC-GTACTCAACTGCAT   1648

Dcluster2       1641 AGGAGCTCACCAGGGAGCAGGCTGGGAGGGGTTCATGCAGAGGGAGCAGC   1690
                     ||||||||.||||||||||||||||||||||||||||||||||.||||||
Dcluster3       1649 AGGAGCTCGCCAGGGAGCAGGCTGGGAGGGGTTCATGCAGAGGCAGCAGC   1698

Dcluster2       1691 TCCCAGCATGCTGCCAGACCCGAGGTAGGGGAGTCCGGATCCAGGCCCGG   1740
                     ||||||||||||||||||||||||||||||||||||||||||||||||||
Dcluster3       1699 TCCCAGCATGCTGCCAGACCCGAGGTAGGGGAGTCCGGATCCAGGCCCGG   1748

Dcluster2       1741 CCTGAAGGGGCTCGGGGGGCCTCTCACATGTAGCAGGGGTGTGCAGGTCC   1790
                     ||||||||||||||||||||||||||||||||||||||||||||||||||
Dcluster3       1749 CCTGAAGGGGCTCGGGGGGCCTCTCACATGTAGCAGGGGTGTGCAGGTCC   1798

Dcluster2       1791 CTTGTGCAGTGTCCAGAGATCATGGACAGGGCGGGCACCGCACCAGAAGC   1840
                     ||||||||||||||||||||||||||||||||||||||.|||||||||||
Dcluster3       1799 CTTGTGCAGTGTCCAGAGATCATGGACAGGGCGGGCACTGCACCAGAAGC   1848

Dcluster2       1841 CAAGCCAGTGACCAGCCTCTGCAGGAAGCACCCATGGGAGCCAACACCTA   1890
                     |||||||||||||||||||||||||||||.|||.||||||||||||||||
Dcluster3       1849 CAAGCCAGTGACCAGCCTCTGCAGGAAGCGCCCGTGGGAGCCAACACCTA   1898

Dcluster2       1891 AGTCTCCCTCCCAGCCCTGCAGCGTGGCCACCCACCACCCCACAGGTCAG   1940
                     ||||||||||||||||||||||||||||||||||||||||||||||||||
Dcluster3       1899 AGTCTCCCTCCCAGCCCTGCAGCGTGGCCACCCACCACCCCACAGGTCAG   1948

Dcluster2       1941 CCTGGGGAGCTCAAGACGGACTGGCGTCCCTACCTGCCAGGTTCCTGTGC   1990
                     ||||||||||||||||||||||||||||||||||||||||||||||||||
Dcluster3       1949 CCTGGGGAGCTCAAGACGGACTGGCGTCCCTACCTGCCAGGTTCCTGTGC   1998

Dcluster2       1991 AGGATGCTGTCCACGGCCGCAATGCTTTGGCCTCTGCTCAGCTTCTGAGG   2040
                     ||||.|||||||||.|.||||||||||.|||||||||||||||||.||||
Dcluster3       1999 AGGACGCTGTCCACAGGCGCAATGCTTGGGCCTCTGCTCAGCTTCCGAGG   2048

Dcluster2       2041 TTCTCCGGCCCACCACCACTGACCTGAATGGGCTTGGAGTGCAGGCCCAG   2090
                     |||.|.|||||||||||||.||||||||.|||||.|||||||||||||||
Dcluster3       2049 TTCCCTGGCCCACCACCACCGACCTGAAGGGGCTCGGAGTGCAGGCCCAG   2098

Dcluster2       2091 CTGAACATCAGACCTGGGCTCAGAGAGCCACTCGGTCCAGGGTGCACTCA   2140
                     |.|.||||.|||||||||||||||.|||||||||||||||||||||||||
Dcluster3       2099 CAGGACATAAGACCTGGGCTCAGATAGCCACTCGGTCCAGGGTGCACTCA   2148

Dcluster2       2141 GCCGTGCACCTGGAGGAGGGTCCTCCATG-CCGCAGGGTTGGCTGGGCAA   2189
                     |||.||||||||||||.|||||||||||| ||||||..||||||||||||
Dcluster3       2149 GCCCTGCACCTGGAGGTGGGTCCTCCATGGCCGCAGCATTGGCTGGGCAA   2198

Dcluster2       2190 GGAGGCATCACAGACCACCCTGCTGTTGGGACGTCCAGCCCTGGACCAGG   2239
                     |||||||||.|||.|||||||||.||||||.|||||||||||||||||||
Dcluster3       2199 GGAGGCATCCCAGGCCACCCTGCCGTTGGGCCGTCCAGCCCTGGACCAGG   2248

Dcluster2       2240 ACTAGGTTCTCAGAATCCTCCCCACCACGGGACCTGTCCATACCTGGACC   2289
                     |||||||||||||||.||||||||||||||||||.|||||.||||||.||
Dcluster3       2249 ACTAGGTTCTCAGAACCCTCCCCACCACGGGACCCGTCCAGACCTGGGCC   2298

Dcluster2       2290 AGCTGACAGCTCCCAAGGGCTCCA-ACCCAGCAGACAAGACCAAGGG--C   2336
                     |||||||||||||||||||||||| ||||||||||||||||||||||  |
Dcluster3       2299 AGCTGACAGCTCCCAAGGGCTCCACACCCAGCAGACAAGACCAAGGGGGC   2348

Dcluster2       2337 GCTGAAGGCGAGCAGCTAGggtttctgatgccggctgtgttgtggtgATG   2386
                     ||||||..|.|.||.||.||||||||||||||||||||||...||||.|.
Dcluster3       2349 GCTGAAAACCAACAACTGGggtttctgatgccggctgtgtcacggtgGTA   2398

Dcluster2       2387 ATACGATAGGTGTGGTTGTAGTTATTGTAGTGTTGCTACcacagtgatgc   2436
                     .|..|.|||.|.|.||.|||||||||.      ||.|||||||||||..|
Dcluster3       2399 TTGTGGTAGCTATTGTGGTAGTTATTA------TGGTACcacagtgacac   2442

Dcluster2       2437 tctcagtgtcagaaaccCCTGCCCCGAGGTCCCAGACCAGGGAGCCTGTC   2486
                     |.||...|.||||||||||.|.|||.||||||||||||||||||||||||
Dcluster3       2443 tgtccaggacagaaaccCCCGTCCCAAGGTCCCAGACCAGGGAGCCTGTC   2492

Dcluster2       2487 CTCCAGGTACGGGGCACCGCCCTGCCCTCGTGACCATCAGACTGCACCCT   2536
                     .||||.|..|||||.||||||||||||||.||||||||||||||||||||
Dcluster3       2493 TTCCAAGACCGGGGTACCGCCCTGCCCTCCTGACCATCAGACTGCACCCT   2542

Dcluster2       2537 GCACGCCCTCACCCTGCTGTCTCAGGGCAGCAACAGCCTCTGGAGGAGTC   2586
                     ||||||||||||||..|||.|||||||.|||||||||||..|||.|.|||
Dcluster3       2543 GCACGCCCTCACCCCACTGGCTCAGGGAAGCAACAGCCTGAGGAAGGGTC   2592

Dcluster2       2587 CTGGGCCAGGCACTCGGCTGTGGAGTCCCAGACAATGATAGCCTCAGCAG   2636
                     |||||||||||.||.||||.|||.|.||||||||||||.|.||.||||||
Dcluster3       2593 CTGGGCCAGGCGCTGGGCTATGGGGACCCAGACAATGAGACCCCCAGCAG   2642

Dcluster2       2637 TTTCCAGCCACCTGACCCTGGAGGAGCTTGTGCAAGCTGGAGGCCGGCTG   2686
                     .||||||||.|||||||||||||||.|||....|.||..|.|.|.|||||
Dcluster3       2643 GTTCCAGCCCCCTGACCCTGGAGGAACTTCCAGAGGCCTGTGCCAGGCTG   2692

Dcluster2       2687 GGGCCCGGAGCCTCAGAACCCTCCCTGCCAGGGGGCCCGTCCAGACCTGG   2736
                     |||.||| |.|||.|||||||||||. |.|..||||||.||||||
Dcluster3       2693 GGGACCG-ACCCTAAGAACCCTCCCC-CGACAGGGCCCATCCAGA-----   2735

Dcluster2       2737 ACCAGCTGACAGCTCCCGAGGACTGCACGCCCACCAGACGAGACCAAGGG   2786
                              ||||||.|||||.||.||.|.|||||||||.||||||||||
Dcluster3       2736 ---------CAGCTCTCGAGGTCTCCATGTCCACCAGACAAGACCAAGGG   2776

Dcluster2       2787 GGCGCTGAAGGCCAGCAGCTGGggtttctgatgccggctgtgtcacggtg   2836
                     |||||||||||||||||.|||||||||.||||||||||||||||||||||
Dcluster3       2777 GGCGCTGAAGGCCAGCAACTGGggtttttgatgccggctgtgtcacggtg   2826

Dcluster2       2837 GTAGTTGTCCTGATGGTTATAGTTATGGTTATGGTTGTGGTTATGGTTAT   2886
                     ||||||
Dcluster3       2827 GTAGTT--------------------------------------------   2832

Dcluster2       2887 GGTTGTAGTGGTTATGATTGTTATGGTTATGGTGGTTATGGTGGTTATGG   2936
                                  |||.|.|||||||||||||||||||||||.|||||||
Dcluster3       2833 -------------ATGGTGGTTATGGTTATGGTGGTTATGGTTGTTATGG   2869

Dcluster2       2937 TGGTTATGGTTATAGTAGTTATAGTTATAGTTATACTTACGAATATACca   2986
                     |                        |||.|||||..|||.|..|||||||
Dcluster3       2870 T------------------------TATGGTTATGGTTATGGTTATACca   2895

Dcluster2       2987 cagtgatactctctgggacaaaaaccCCTGCCCCTGAGGGTCCACGGCCA   3036
                     ||||||||||||||||||||||||||||||||||||||||||||||.|||
Dcluster3       2896 cagtgatactctctgggacaaaaaccCCTGCCCCTGAGGGTCCACGACCA   2945

Dcluster2       3037 GGGATCCTGGAGGCTGTGTTTGGGGCACCGCCCTGCCCTCCTGCCATTGG   3086
                     ||||..|||.|.|||||||||||||||||.||||||||||||||||||||
Dcluster3       2946 GGGAATCTGTAAGCTGTGTTTGGGGCACCACCCTGCCCTCCTGCCATTGG   2995

Dcluster2       3087 ACTACAACCTGCACGCCCTCACCCTGCTGACTCGGGCAGCAACAGCCTAA   3136
                     |||.||.|||||||||||||||||||||||||||||||||||||||||||
Dcluster3       2996 ACTGCACCCTGCACGCCCTCACCCTGCTGACTCGGGCAGCAACAGCCTAA   3045

Dcluster2       3137 GATAAGAGTCCTGGGCCAGGCGCTGGGCTGTGGATACCCAGACAATGAGA   3186
                     ||||||.|||||||||||||.|||||||||||||||||||||||||||||
Dcluster3       3046 GATAAGGGTCCTGGGCCAGGAGCTGGGCTGTGGATACCCAGACAATGAGA   3095

Dcluster2       3187 TCTCCAGCAGGTTCCCACCTCCTGACCCTGGAGGAACTTCCGCAGCCCTG   3236
                     ||||||||||||||||.||||||||.| ||||||||||.||||||.||||
Dcluster3       3096 TCTCCAGCAGGTTCCCGCCTCCTGATC-TGGAGGAACTGCCGCAGGCCTG   3144

Dcluster2       3237 TGGCAGGCTGGGGACCGACCCTAAGAACCCTCCCCCC-------------   3273
                     ||.||||||||||||||||||||||||||||||||||
Dcluster3       3145 TGCCAGGCTGGGGACCGACCCTAAGAACCCTCCCCCCCGCAGGGCCATCC   3194

Dcluster2       3274 --------------------------------------------------   3273

Dcluster3       3195 AGACCTGGGCCAGCTGACAGCTCCCGAGGACTCCACGTCCACCAGACGAG   3244

Dcluster2       3274 --------------------------------------------------   3273

Dcluster3       3245 ACCAAGGGGACGCCGAAGGCCAGCAGCTGGggtttctgatgccggctgtg   3294

Dcluster2       3274 --------------------------------------------------   3273

Dcluster3       3295 ttgtggtgATGATACGATAGGTGTGGTTTTAGTTATTGTAGTGTTGCTAC   3344

Dcluster2       3274 --------------------------------------------------   3273

Dcluster3       3345 cacagtgacgctctcagtgtcagaaaccCCTGCCCCGAGGTCCCAGACCA   3394

Dcluster2       3274 --------------------------------------------------   3273

Dcluster3       3395 GGGAGCTTGTCCTCCAGGTCCGGGGTACGGCCCTGCCCTCCTGACCATCG   3444

Dcluster2       3274 --------------------------------------------------   3273

Dcluster3       3445 GACTGCACCCTGCATGCCCTCACCCTGCTGACTCAGGGCAGCAACAGAAT   3494

Dcluster2       3274 --------------------------------------------------   3273

Dcluster3       3495 GAGGAAGGGTCCTGGGCCAAGAGCTTGGCTGCTGGGACCCAGACAATGAG   3544

Dcluster2       3274 --------------------------------------------------   3273

Dcluster3       3545 GCCCCCAGCAAGTTCCAGCTCCCTGACCCTGGAGGAGGTTGTGCAGGTCA   3594

Dcluster2       3274 ----------------------------------------GCAGGGCCCG   3283
                                                             ||||||||||
Dcluster3       3595 GAGGCCGGCTGGGGACCGGATCCTCAGAACCCTCCCTGCCGCAGGGCCCG   3644

Dcluster2       3284 TCCAGACCTGAACCAGCTGACAGCTCCCGAGGGCTCCACGCCCCCCAGAC   3333
                     ||||||| ||.|||||||||||||||||.|||.||||||||||.||||||
Dcluster3       3645 TCCAGAC-TGGACCAGCTGACAGCTCCCAAGGACTCCACGCCCACCAGAC   3693

Dcluster2       3334 GAGACCAAGGGGGCGCCGAAGGCCAGCAGCTGGggtttctgatgccggct   3383
                     ||||||||||||||||||||||||||||||.|||||||||||||||.|||
Dcluster3       3694 GAGACCAAGGGGGCGCCGAAGGCCAGCAGCCGGggtttctgatgccagct   3743

Dcluster2       3384 gtgtcacggtgGTAGTTGTTATAGTGGTTATGGTTATGGTTGTGGTTATG   3433
                     |||||||||||||||||||||||||||||||||||||||||.|||||.||
Dcluster3       3744 gtgtcacggtgGTAGTTGTTATAGTGGTTATGGTTATGGTTATGGTTGTG   3793

Dcluster2       3434 GTTATGGTTATGATTATACcacagtgacactctctgggacaaaaacc   3480
                     ||||||||||||.||||||||||||||||||||||||||||||||||
Dcluster3       3794 GTTATGGTTATGGTTATACcacagtgacactctctgggacaaaaacc   3840
```
