## Supplemental Fig. 3 for "Formation of ultralong DH regions through genomic rearrangement"

**Supplemental Figure 3.** Homology of IGHD regions from clusters 2 and 3. The DH regions IGHD1-2, IGHD2-2, IGHD5-2, IGHD8-2, and IGHD6-2 from cluster 2 and IGHD1-3, IGHD2-3, IGHD3-3, IGHD7-3, IGHD5-3, and IGHD6-3 from cluster 3 were aligned using Clustal2.1 and phylogenetic tree (A) and percent identity matrix (B) shown. The IGHD3-3 or IGHD7-3 regions do not have clear paralogs on cluster 2. Paralogs on the two clusters are highlighted green. Percent identities above 95% are in bold.

**A.**

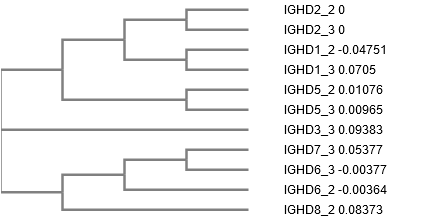

**B.**

```
#  Percent Identity  Matrix - created by Clustal2.1
#
#
		2-2     2-3     1-2     1-3     5-2      5-3    3-3      7-3     8-2     6-2     6-3
     1: IGHD2_2     100.00  100.00   43.75   36.11   48.61   47.22   45.83   48.61   48.61   50.00   48.33
     2: IGHD2_3     100.00  100.00   43.75   36.11   48.61   47.22   45.83   48.61   48.61   50.00   48.33
     3: IGHD1_2      43.75   43.75  100.00   97.70   55.17   55.17   58.06   44.12   47.06   60.00   57.78
     4: IGHD1_3      36.11   36.11   97.70  100.00   39.80   40.82   43.48   32.79   40.46   47.19   46.07
     5: IGHD5_2      48.61   48.61   55.17   39.80  100.00   97.96   71.11   75.29   72.45   78.08   76.71
     6: IGHD5_3      47.22   47.22   55.17   40.82   97.96  100.00   72.22   74.12   71.43   79.45   78.08
     7: IGHD3_3      45.83   45.83   58.06   43.48   71.11   72.22  100.00   76.62   80.43   88.00   85.33
     8: IGHD7_3      48.61   48.61   44.12   32.79   75.29   74.12   76.62  100.00   81.30   95.00   95.00
     9: IGHD8_2      48.61   48.61   47.06   40.46   72.45   71.43   80.43   81.30  100.00   89.47   89.47
    10: IGHD6_2      50.00   50.00   60.00   47.19   78.08   79.45   88.00   95.00   89.47  100.00   96.49
    11: IGHD6_3      48.33   48.33   57.78   46.07   76.71   78.08   85.33   95.00   89.47   96.49  100.00
```
