## Supplemental Fig. 4 for "Formation of ultralong DH regions through genomic rearrangement"

**Supplemental Figure 4.** Homology of intergenic regions. The intergenic regions between IGHD2-2 and IGHD5-2 (before5-2), IGHD5-2 and IGHD8-2 (before8-2), IGHD8-2 and IGHD6-2 (before6-2) of cluster 2 and IGHD2-3 and IGHD3-3 (before3-3), IGHD3-3 and IGHD7-3 (before7-3), IGHD7-3 and IGHD5-3 (before 5-3), and IGHD5-3 and IGHD6-3 (before6-3) of cluster 3 were aligned using Clustal2.1 and phylogenetic tree (A) and percent identity matrix (B) shown. The region between IGHD3-3 and IGHD7-3 (before7-3) does not have a clear paralog on cluster 3. Paralogs on the two clusters are highlighted green.

**A.**

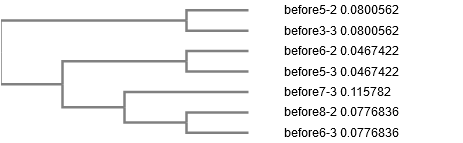

**B.**

```
#  Percent Identity  Matrix - created by Clustal2.1
#
#
			b5-2	b3-3	b6-2	b5-3	b7-3	b8-2	b6-3
     1: before5-2   100.00   85.23   44.17   42.46   43.27   48.78   49.85
     2: before3-3    85.23  100.00   52.45   51.38   51.92   57.32   57.19
     3: before6-2    44.17   52.45  100.00   92.35   76.19   74.15   73.50
     4: before5-3    42.46   51.38   92.35  100.00   77.61   73.22   72.57
     5: before7-3    43.27   51.92   76.19   77.61  100.00   80.53   80.24
     6: before8-2    48.78   57.32   74.15   73.22   80.53  100.00   87.29
     7: before6-3    49.85   57.19   73.50   72.57   80.24   87.29  100.00
```
