## Supplemental Fig. 5 for "Formation of ultralong DH regions through genomic rearrangement"

**Supplemental Figure 5.** Local alignment of IGHD7-3 and IGHD7-4 with IGHD8-2.

```
########################################
# Program: matcher
# Rundate: Sat  5 Oct 2019 00:28:58
# Commandline: matcher
#    -auto
#    -stdout
#    -asequence emboss_matcher-I20191005-002855-0888-87948780-p2m.asequence
#    -bsequence emboss_matcher-I20191005-002855-0888-87948780-p2m.bsequence
#    -datafile EDNAFULL
#    -gapopen 16
#    -gapextend 4
#    -alternatives 1
#    -aformat3 pair
#    -snucleotide1
#    -snucleotide2
# Align_format: pair
# Report_file: stdout
########################################

#=======================================
#
# Aligned_sequences: 2
# 1: IGHD8_2
# 2: IGHD7_3
# Matrix: EDNAFULL
# Gap_penalty: 16
# Extend_penalty: 4
#
# Length: 97
# Identity:      83/97 (85.6%)
# Similarity:    83/97 (85.6%)
# Gaps:           0/97 ( 0.0%)
# Score: 359
#
#
#=======================================

IGHD8_2          108 TGGTGGTTATGGTGGTTATGGTGGTTATGGTTATAGTAGTTATAGTTATA    157
                     ||||.|||||||||||||||||..|..|||||||.||.|||||.|||||.
IGHD7_3           27 TGGTAGTTATGGTGGTTATGGTTATGGTGGTTATGGTTGTTATGGTTATG     76

IGHD8_2          158 GTTATACTTACGAATATACCACAGTGATACTCTCTGGGACAAAAACC    204
                     |||||..|||.|..|||||||||||||||||||||||||||||||||
IGHD7_3           77 GTTATGGTTATGGTTATACCACAGTGATACTCTCTGGGACAAAAACC    123


#---------------------------------------
#---------------------------------------


########################################
# Program: matcher
# Rundate: Sat  5 Oct 2019 00:30:17
# Commandline: matcher
#    -auto
#    -stdout
#    -asequence emboss_matcher-I20191005-003015-0194-38128626-p2m.asequence
#    -bsequence emboss_matcher-I20191005-003015-0194-38128626-p2m.bsequence
#    -datafile EDNAFULL
#    -gapopen 16
#    -gapextend 4
#    -alternatives 1
#    -aformat3 pair
#    -snucleotide1
#    -snucleotide2
# Align_format: pair
# Report_file: stdout
########################################

#=======================================
#
# Aligned_sequences: 2
# 1: IGHD8_2
# 2: IGHD7_4
# Matrix: EDNAFULL
# Gap_penalty: 16
# Extend_penalty: 4
#
# Length: 132
# Identity:     102/132 (77.3%)
# Similarity:   102/132 (77.3%)
# Gaps:           3/132 ( 2.3%)
# Score: 378
#
#
#=======================================

IGHD8_2           73 GGTTATGGTTGTAGTGGTTATGATTGTTATGGTTATGGTGGTTATGGTGG    122
                     ||||.|.|.||..|....|.|.|..||..|.|||||||||||||||   |
IGHD7_4            1 GGTTTTTGATGCCGGCTGTGTCACGGTGGTAGTTATGGTGGTTATG---G     47

IGHD8_2          123 TTATGGTGGTTATGGTTATAGTAGTTATAGTTATAGTTATACTTACGAAT    172
                     |||||||||||||||||.|..|.|||||.|||||.|||||..|||.|..|
IGHD7_4           48 TTATGGTGGTTATGGTTGTTATGGTTATGGTTATGGTTATGGTTATGGTT     97

IGHD8_2          173 ATACCACAGTGATACTCTCTGGGACAAAAACC    204
                     ||||||||||||||||||||||||||||||||
IGHD7_4           98 ATACCACAGTGATACTCTCTGGGACAAAAACC    129


#---------------------------------------
#---------------------------------------
```
