## Supplemental Fig. 6 for "Formation of ultralong DH regions through genomic rearrangement"

**Supplemental Figure 6.** Ultralong DH region variants differ in in length. A multiple sequence alignment was performed using MUSCLE with the two *Bos taurus* IGHD8-2 polymorphisms, as well as IGHD8-2 homologs from *Bison bison* and *Bos grunniens*. The RSS are in lowercase, with heptamer and nonamer underlined and in italics.

```
CLUSTAL multiple sequence alignment by MUSCLE (3.8)


IGHD8-2*01                            ggtttctgatgccggctgtgtcacggtgGTAGTTGTCCTGATGGTTATAGTTAT------
IGHD8-2*02                            ggtttctgatgccggctgtgtcacggtgGTAGTTGTCCTGATGGTTATAGTTAT------
Bison_IGHD8-2                         ggtttctgatgccggctgtgtcacggtgGTAGTTGTCCTGATGGTTATAGTTATGGTTAT
BosGru_IGHD8-2                        ggtttctgatgccggctgtgtcacggtgGTAGTTGTCCTGATGGTTATGGTTATGGTTAT
                                      ************************************************ *****

IGHD8-2*01                            GGTTATGGTTGTGGTTATG---GTTATGGTTGTAGTGGTTATGATTGTTATGGTTATGGT
IGHD8-2*02                            GGTTATGGTTGTGGTTATGGTAGTTATGGTTGTAGTGGTTATGATTGTTATGGTTATGGT
Bison_IGHD8-2                         GGTTATGGTTGTGGTTATGGTGGTTATGGTTGTAGTGGTTATGATTGTTATGGTTATGGT
BosGru_IGHD8-2                        GGTTATGGTTGTGGTTATGGTGGTTATGGTTGTAGTGGTTATGATTGTTATGGTTATGGT
                                      *******************   **************************************

IGHD8-2*01                            GGTTATGGTGGTTATGGTGGTTA---TGGTTATAGTAGTTATAGTTATAGTTATACTTAC
IGHD8-2*02                            GGTTATGGTGGTTATGGTGGTTATGGTGGTTATAGTAGTTATAGTTATAGTTATAGTTAC
Bison_IGHD8-2                         AGTTA---------------------TGGTTATAGTGGTTATAGTTATAGTTATAGTTAC
BosGru_IGHD8-2                        GGTTA---------------------TGGTTATAGTGGTTATAGTTATAGTTATAGTTAC
                                       ****                     ********** ****************** ****

IGHD8-2*01                            GAATATACcacagtgatactctctgggacaaaaacc
IGHD8-2*02                            GAATATACcacagtgatactctctgggacaaaaacc
Bison_IGHD8-2                         GAATATACcacagtgatactctctgggacaaaaacc
BosGru_IGHD8-2                        GAATATACcacagtgatactctctgggacaaaaacc
                                      ************************************
```
