## Supplemental Fig. 7 for "Formation of ultralong DH regions through genomic rearrangement"

**Supplemental Figure 7.** Alignment of IGHD3 and IGHD6 family members with IGHD8-2.

```
########################################
# Program: matcher
# Rundate: Sat  5 Oct 2019 00:41:09
# Commandline: matcher
#    -auto
#    -stdout
#    -asequence emboss_matcher-I20191005-004107-0321-40057744-p1m.asequence
#    -bsequence emboss_matcher-I20191005-004107-0321-40057744-p1m.bsequence
#    -datafile EDNAFULL
#    -gapopen 16
#    -gapextend 4
#    -alternatives 1
#    -aformat3 pair
#    -snucleotide1
#    -snucleotide2
# Align_format: pair
# Report_file: stdout
########################################

#=======================================
#
# Aligned_sequences: 2
# 1: IGHD8_2
# 2: IGHD3_4
# Matrix: EDNAFULL
# Gap_penalty: 16
# Extend_penalty: 4
#
# Length: 65
# Identity:      52/65 (80.0%)
# Similarity:    52/65 (80.0%)
# Gaps:           5/65 ( 7.7%)
# Score: 184
#
#
#=======================================

IGHD8_2            1 GGTTTCTGATGCCGGCTGTGTCACGGTGGTA--GTTGTCCTGATGGTTAT     48
                     |||||||||||||||||||||||||||||||  ||.||....||.||..|
IGHD3_4            1 GGTTTCTGATGCCGGCTGTGTCACGGTGGTATTGTGGTAGCTATTGTGGT     50

IGHD8_2           49 AGTTATGGTTATGGT     63
                     ||||||   ||||||
IGHD3_4           51 AGTTAT---TATGGT     62


#---------------------------------------
#---------------------------------------


########################################
# Program: matcher
# Rundate: Sat  5 Oct 2019 00:42:25
# Commandline: matcher
#    -auto
#    -stdout
#    -asequence emboss_matcher-I20191005-004222-0822-86095519-p2m.asequence
#    -bsequence emboss_matcher-I20191005-004222-0822-86095519-p2m.bsequence
#    -datafile EDNAFULL
#    -gapopen 16
#    -gapextend 4
#    -alternatives 1
#    -aformat3 pair
#    -snucleotide1
#    -snucleotide2
# Align_format: pair
# Report_file: stdout
########################################

#=======================================
#
# Aligned_sequences: 2
# 1: IGHD8_2
# 2: IGHD3_3
# Matrix: EDNAFULL
# Gap_penalty: 16
# Extend_penalty: 4
#
# Length: 65
# Identity:      52/65 (80.0%)
# Similarity:    52/65 (80.0%)
# Gaps:           5/65 ( 7.7%)
# Score: 184
#
#
#=======================================

IGHD8_2            1 GGTTTCTGATGCCGGCTGTGTCACGGTGGTA--GTTGTCCTGATGGTTAT     48
                     |||||||||||||||||||||||||||||||  ||.||....||.||..|
IGHD3_3            1 GGTTTCTGATGCCGGCTGTGTCACGGTGGTATTGTGGTAGCTATTGTGGT     50

IGHD8_2           49 AGTTATGGTTATGGT     63
                     ||||||   ||||||
IGHD3_3           51 AGTTAT---TATGGT     62


#---------------------------------------
#---------------------------------------


########################################
# Program: matcher
# Rundate: Fri  4 Oct 2019 23:30:39
# Commandline: matcher
#    -auto
#    -stdout
#    -asequence emboss_matcher-I20191004-233038-0324-2606874-p2m.asequence
#    -bsequence emboss_matcher-I20191004-233038-0324-2606874-p2m.bsequence
#    -datafile EDNAFULL
#    -gapopen 16
#    -gapextend 4
#    -alternatives 1
#    -aformat3 pair
#    -snucleotide1
#    -snucleotide2
# Align_format: pair
# Report_file: stdout
########################################

#=======================================
#
# Aligned_sequences: 2
# 1: IGHD8_2
# 2: IGHD6_2
# Matrix: EDNAFULL
# Gap_penalty: 16
# Extend_penalty: 4
#
# Length: 85
# Identity:      76/85 (89.4%)
# Similarity:    76/85 (89.4%)
# Gaps:           0/85 ( 0.0%)
# Score: 344
#
#
#=======================================

IGHD8_2            1 GGTTTCTGATGCCGGCTGTGTCACGGTGGTAGTTGTCCTGATGGTTATAG     50
                     ||||||||||||||||||||||||||||||||||||..|..|||||||.|
IGHD6_2            1 GGTTTCTGATGCCGGCTGTGTCACGGTGGTAGTTGTTATAGTGGTTATGG     50

IGHD8_2           51 TTATGGTTATGGTTGTGGTTATGGTTATGGTTGTA     85
                     ||||||||.|||||.||||||||||||||.||.||
IGHD6_2           51 TTATGGTTGTGGTTATGGTTATGGTTATGATTATA     85


#---------------------------------------
#---------------------------------------


########################################
# Program: matcher
# Rundate: Fri  4 Oct 2019 23:33:07
# Commandline: matcher
#    -auto
#    -stdout
#    -asequence emboss_matcher-I20191004-233305-0188-59118204-p1m.asequence
#    -bsequence emboss_matcher-I20191004-233305-0188-59118204-p1m.bsequence
#    -datafile EDNAFULL
#    -gapopen 16
#    -gapextend 4
#    -alternatives 1
#    -aformat3 pair
#    -snucleotide1
#    -snucleotide2
# Align_format: pair
# Report_file: stdout
########################################

#=======================================
#
# Aligned_sequences: 2
# 1: IGHD8_2
# 2: IGHD6_3
# Matrix: EDNAFULL
# Gap_penalty: 16
# Extend_penalty: 4
#
# Length: 85
# Identity:      78/85 (91.8%)
# Similarity:    78/85 (91.8%)
# Gaps:           0/85 ( 0.0%)
# Score: 362
#
#
#=======================================

IGHD8_2            1 GGTTTCTGATGCCGGCTGTGTCACGGTGGTAGTTGTCCTGATGGTTATAG     50
                     |||||||||||||.||||||||||||||||||||||..|..|||||||.|
IGHD6_3            1 GGTTTCTGATGCCAGCTGTGTCACGGTGGTAGTTGTTATAGTGGTTATGG     50

IGHD8_2           51 TTATGGTTATGGTTGTGGTTATGGTTATGGTTGTA     85
                     ||||||||||||||||||||||||||||||||.||
IGHD6_3           51 TTATGGTTATGGTTGTGGTTATGGTTATGGTTATA     85


#---------------------------------------
#---------------------------------------
```
