## Supplemental Fig. 8 for "Formation of ultralong DH regions through genomic rearrangement"

**Supplemental Figure 8.** Fusion between DH regions to create a hybrid ultralong DH. IGHD6-3 and IGHD7-3 were fused to create an ultralong DH region. Translated amino acids are above the sequence. Red nucleotides are a TGGTTA repeat, blue are TAGTTA, and green are TGGTTG. The RSS are in lower case with heptamer and nonamer in italics and underlined. Note that the repeating units encode amino acid repeats, and that the differences in length between the artificial hybrid and natural IGHD8-2 regions can be attributed to the number of repeats.

IGHD6_3

*ggtttctga*tgccagctgtgt*cacggtg*GTAGTTGTTATAGTGGTTATGGTTATGGTTATGGTTGTGGTTATGGTTATGGTTATAC*cacagtg*acactctctggg*acaaaaacc*

>IGHD7_3

*Ggtttttga*tgccggctgtgt*cacggtg*GTAGTTATGGTGGTTATGGTTATGGTGGTTATGGTTGTTATGGTTATGGTTATGGTTATGGTTATAC*cacagtg*atactctctggg*acaaaaacc*

Hybrid	S  C  Y  S  G  Y  G  Y  G  Y  G  C  G  Y  G  Y  G  Y  S  S  Y  G  G  Y  G  Y  G  G  Y  G  C  Y  G  Y  G  Y  G  Y  G  Y

*ggtttctga*tgccagctgtgt*cacggtg*GTAGTTGTTATAGTGGTTATGGTTATGGTTATGGTTGTGGTTATGGTTATGGTTATAGTAGTTATGGTGGTTATGGTTATGGTGGTTATGGTTGTTATGGTTATGGTTATGGTTATGGTTATAC*cacagtg*atactctctggg*acaaaaacc*

Bison IGHD8-2   S  C  P  D  G  Y  S  Y  G  Y  G  Y  G  C  G  Y  G  G  Y  G  C  S  G  Y  D  C  Y  G  Y  G  S  Y  G  Y  S  G  Y  S  Y  S  Y  S  Y  E  Y

ggtttctgatgccggctgtgtcacggtgGTAGTTGTCCTGATGGTTATAGTTATGGTTATGGTTATGGTTGTGGTTATGGTGGTTATGGTTGTAGTGGTTATGATTGTTATGGTTATGGTAGTTATGGTTATAGTGGTTATAGTTATAGTTATAGTTACGAATATACcacagtgatactctctgggacaaaaacc

IGHD8-2*01  S  C  P  D  G  Y  S  Y  G  Y  G  C  G  Y  G  Y  G  C  S  G  Y  D  C  Y  G  Y  G  G  Y  G  G  Y  G  G  Y  G  Y  S  S  Y  S  Y  S  Y  T  Y  E  Y

ggtttctgatgccggctgtgtcacggtgGTAGTTGTCCTGATGGTTATAGTTATGGTTATGGTTGTGGTTATGGTTATGGTTGTAGTGGTTATGATTGTTATGGTTATGGTGGTTATGGTGGTTATGGTGGTTATGGTTATAGTAGTTATAGTTATAGTTATACTTACGAATATACcacagtgatactctctgggacaaaaacc

IGHD8-2*02	S  C  P  D  G  Y  S  Y  G  Y  G  C  G  Y  G  S  Y  G  C  S  G  Y  D  C  Y  G  Y  G  G  Y  G  G  Y  G  G  Y  G  G  Y  S  S  Y  S  Y  S  Y  S  Y  E  Y

ggtttctgatgccggctgtgtcacggtgGTAGTTGTCCTGATGGTTATAGTTATGGTTATGGTTGTGGTTATGGTAGTTATGGTTGTAGTGGTTATGATTGTTATGGTTATGGTGGTTATGGTGGTTATGGTGGTTATGGTGGTTATAGTAGTTATAGTTATAGTTATAGTTACGAATATACcacagtgatactctctgggacaaaaacc
